# A disordered encounter complex is central to the yeast Abp1p SH3 domain binding pathway

**DOI:** 10.1101/2020.03.23.003269

**Authors:** Gabriella J. Gerlach, Rachel Carrock, Robyn Stix, Elliott J. Stollar, K. Aurelia Ball

**Affiliations:** Skidmore College, Department of Chemistry, 815 N Broadway, Saratoga Springs, NY 12866, United States; University of Liverpool, United Kingdom

## Abstract

Protein-protein interactions are involved in a wide range of cellular processes. These interactions often involve intrinsically disordered proteins (IDPs) and protein binding domains. However, the details of IDP binding pathways are hard to characterize using experimental approaches, which can rarely capture intermediate states present at low populations. SH3 domains are common protein interaction domains that typically bind proline-rich disordered segments and are involved in cell signaling, regulation, and assembly. We hypothesized, given the flexibility of SH3 binding peptides, that their binding pathways include multiple steps important for function. Molecular dynamics simulations were used to characterize the steps of binding between the yeast Abp1p SH3 domain (AbpSH3) and a proline-rich IDP, ArkA. Before binding, the N-terminal segment 1 of ArkA is pre-structured and adopts a polyproline II helix, while segment 2 of ArkA (C-terminal) adopts a 3^10^ helix, but is far less structured than segment 1. As segment 2 interacts with AbpSH3, it becomes more structured, but retains flexibility even in the fully engaged state. Binding simulations reveal that ArkA enters a flexible encounter complex before forming the fully engaged bound complex. In the encounter complex, transient nonspecific hydrophobic and long-range electrostatic contacts form between ArkA and the binding surface of SH3. The encounter complex ensemble includes conformations with segment 1 in both the forward and reverse orientation, suggesting that segment 2 may play a role in stabilizing the correct binding orientation. While the encounter complex forms quickly, the slow step of binding is the transition from the disordered encounter ensemble to the fully engaged state. In this transition, ArkA makes specific contacts with AbpSH3 and buries more hydrophobic surface. Simulating the binding between ApbSH3 and ArkA provides insight into the role of encounter complex intermediates and nonnative hydrophobic interactions for other SH3 domains and IDPs in general.

**Author Summary:** Complex cellular processes are mediated by interactions between proteins, and to determine how these interactions affect cellular function and binding kinetics we often must understand the protein binding pathway. Many protein interaction domains, such as the SH3 domain, bind to intrinsically disordered proteins in a coupled folding and binding process. Using molecular dynamics simulations, we find that the binding of the disordered ArkA peptide to the yeast Abp1p SH3 domain proceeds through a flexible, disordered encounter complex before reaching a stable fully bound state. The encounter complex is stabilized by nonspecific long-range electrostatic interactions and nonspecific hydrophobic interactions between the peptide and domain. Our simulations highlight the important role of hydrophobic interactions in the entire SH3 binding process: both nonspecific hydrophobic contacts in the encounter complex and specific hydrophobic contacts in the fully bound complex. The encounter complex could be key to understanding the functional behavior of SH3 domain interactions because the encounter complex forms very quickly and the transition to the fully bound state is slower. In cells, an SH3 domain may form an encounter complex quickly and nonspecifically with many potential binding partners, allowing it to search for the correct recognition sequence before completing the binding process.

## Introduction

Protein-protein interactions are involved in most cellular processes, especially cellular signaling. These interactions often involve binding of small protein domains to intrinsically disordered proteins (IDP), but unlike long-lived complexes typically involving larger and stronger interfaces, the binding pathways for these interactions are not always well understood [1–3]. Regions of disorder are now known to be present in between 25% and 41% of eukaryotic proteins, and can exhibit functional diversity by having multiple interaction partners [4]. IDPs also tend to bind with lower affinity to their partners than folded proteins, with fast association and dissociation [3,5–7]. This fast binding and unbinding, along with the fast turnover rates of IDPs within cells, allows for regulation of processes that require rapid responses [8]. Despite fast on and off rates, IDP binding interactions must still be very specific in order to relay signals accurately, which may require more complex binding landscapes [3,5,6,9,10]. To fully understand how IDPs bind their partners, how their binding is modulated by different cellular contexts, and how changes to the binding process can be used to regulate function, it is essential to go beyond analyzing the final bound state and instead characterize the complete binding pathway and associated kinetics.

IDPs often bind to folded proteins through a pathway that takes place in at least two steps [11–15]. Binding typically begins with the creation of an encounter complex ensemble when the IDP “dances” on top of its partner domain before transitioning to a more structured fully engaged bound state through a process of induced-fit folding [13,15–19]. IDPs are well suited to quickly form this initial encounter complex because they generally adopt a more extended conformational ensemble and therefore have a larger capture radius than folded proteins of the same length [20]. Electrostatic interactions have been shown to often drive the formation of encounter complex ensembles and can even accelerate association beyond the diffusion limited rate, predominantly by electrostatic orientational steering [5,11,13,21–23]. Additionally, if one segment of the IDP possesses more intrinsic pre-folded structure, binding may proceed starting with this segment and then extending to the rest of the sequence which folds upon interaction with the partner protein [11, 24]. Thus, pre-formed secondary structure can improve affinity and influence the binding pathway, including the nature of intermediate states [25, 26], but too much structure can slow binding without improving affinity [11, 27]. Additionally, the ability of IDPs to form nonnative contacts during binding, and the presence of significant disorder even after binding can also be important for IDP function [20,27,28]. This underscores the importance of understanding the interactions at play during the binding process as well as in the fully engaged final complex.

In addition to the nature of the intermediate states in the binding pathway, the location of the transition state in the pathway dictates the binding kinetics and is therefore critical to function. The transition state for binding can either precede or follow the encounter complex intermediate [5]. Fast-binding proteins are canonically thought to bind in a diffusion-limited manner, and therefore experience a rate determining transition state that precedes the encounter complex [5]. In other cases, when electrostatic attraction enhances binding, binding can proceed completely downhill, without a free energy barrier [29, 30]. However, weaker IDP complexes with short lifetimes may exhibit binding kinetics that are different in nature from higher affinity complexes [31], and a few IDPs have been shown to associate quickly to form an encounter complex followed by a slower transition to the fully engaged complex [14,17,19,32–37]. Because of their fast binding and dissociation rates and short-lived intermediate states, IDP binding often appears as two-state in NMR [38–40] and stopped-flow experiments [41–43]. Due to experimental challenges, for many IDPs the specific binding pathway, including the nature of binding intermediates and the timescale of their formation, is still unknown.

Computer simulations have been a valuable tool for examining the binding pathway at temporal and spatial resolutions that cannot be obtained through experiments. Initially, coarse-grained molecular dynamics (MD) simulations based on the topology of the fully engaged complex indicated that the initial step in the binding process for IDPs might often be the formation of a flexible encounter complex [13,37,44–50]. Another strategy to simulate IDP binding and characterize the encounter complex with limited computational power has been to conduct atomistic MD using an advanced sampling algorithm [51], such as multicanonical MD [52]. More recently, advancements in both hardware and the accuracy of force fields have enabled unbiased atomistic MD simulations of IDP binding on the microsecond timescale [53]. Those unbiased MD simulations that have explicitly examined the IDP binding pathway generally reveal a fast initial association between the IDP and its partner, followed by a slower evolution into the fully engaged complex [23,54–58]. However, the details of the binding pathway, including the nature of intermediates in the binding process, has yet to be determined for most IDPs and their binding partners.

One common IDP binding domain is the SH3 domain. It is conserved through more than one billion years of evolution from yeast to humans, and frequently occurs in protein-protein interaction modules, often involving cellular signaling, assembly, or regulation [59]. SH3 domains bind disordered proline-rich target peptides that usually contain a PxxP motif, where x can be any residue [60, 61]. This PxxP motif forms a polyproline type II (PPII) helix in the bound complex and is flanked by specificity elements, which often include positively charged lysine or arginine residues [59, 62]. The PxxP motif, which is pseudo-palindromic, has been observed to bind the SH3 domain surface in two different orientations (class I and class II) depending on the location of positively charged residues either N or C terminal to the PxxP motif (+xxPxxP or PxxPx+) [63, 64]. Despite very similar binding motifs and bound structures, SH3 domains perform a wide variety of different functions in different contexts that require specific binding interactions and biophysical properties [65–67]. Understanding the SH3 domain-peptide binding process may help to reveal the mechanism for the functional diversity of SH3 domains and serve as a model for understanding binding properties of extended IDPs.

Previous studies of proline-rich peptides binding to SH3 domains indicated that fully engaged bound complexes often exhibit conformational exchange between different bound states [39,40,68]. However, information about the binding pathway is more limited, as NMR experiments on SH3 domains often indicate two-state binding, possibly due to fast exchange of an encounter complex with either the fully engaged bound state or the unbound state [23,38–40]. One study of SH3-peptide binding found that the transition state for binding is stabilized by long-range electrostatic interactions; however, there is less electrostatic enhancement to the binding rate than for folded proteins, which form more short-range electrostatic interactions in the transition state [31]. Simulation studies of the C-CRK N-terminal SH3 domain binding to a proline-rich peptide have also indicated that electrostatic interactions are important for the formation of the highly dynamic encounter complex, which transitions to the fully engaged complex when the PPII helix locks into the hydrophobic grooves of the binding site [21,23,56]. However, these results are somewhat in contrast to the picture that hydrophobic interactions are most important for stabilizing the encounter complex for IDPs [17], and the authors of these studies did not try to quantitatively show that an encounter complex is an intermediate to binding or assess the different types of intermolecular interactions in the encounter complex across many independent binding simulations [23,56,69]. Therefore, it is still not clear that SH3 domains form a metastable electrostatic encounter complex, or whether the transition state for SH3 binding occurs before or after the formation of such an encounter complex.

We have used all-atom molecular dynamics (MD) simulations to characterize the initial binding interaction between an SH3 domain of yeast Actin Binding Protein 1 (Abp1p) and ArkA, a disordered region of the yeast actin patch kinase, Ark1p. Abp1p is involved in assembly of the actin cytoskeleton through localization of cortical actin patches, actin organization, and endocytosis [70, 71]. While several other sequences are known to bind the Abp1p SH3 domain (AbpSH3), ArkA is the partner with the highest affinity for the domain [72]. The structures of AbpSH3 alone and bound to ArkA (Fig 1A) have been solved by x-ray crystallography and NMR, respectively [70, 72]. We focus on the binding of a 12-residue truncation of ArkA (residues Lys(3) to Lys(−8)) that binds AbpSH3 with a *K*_d_ of 1.7 μM and is comprised of an N-terminal segment containing the PxxP motif and an adjacent C-terminal segment containing key specificity elements (we use a standard numbering system for peptide positions based on [63], as shown in Fig 1B) [72].

**Fig 1.**
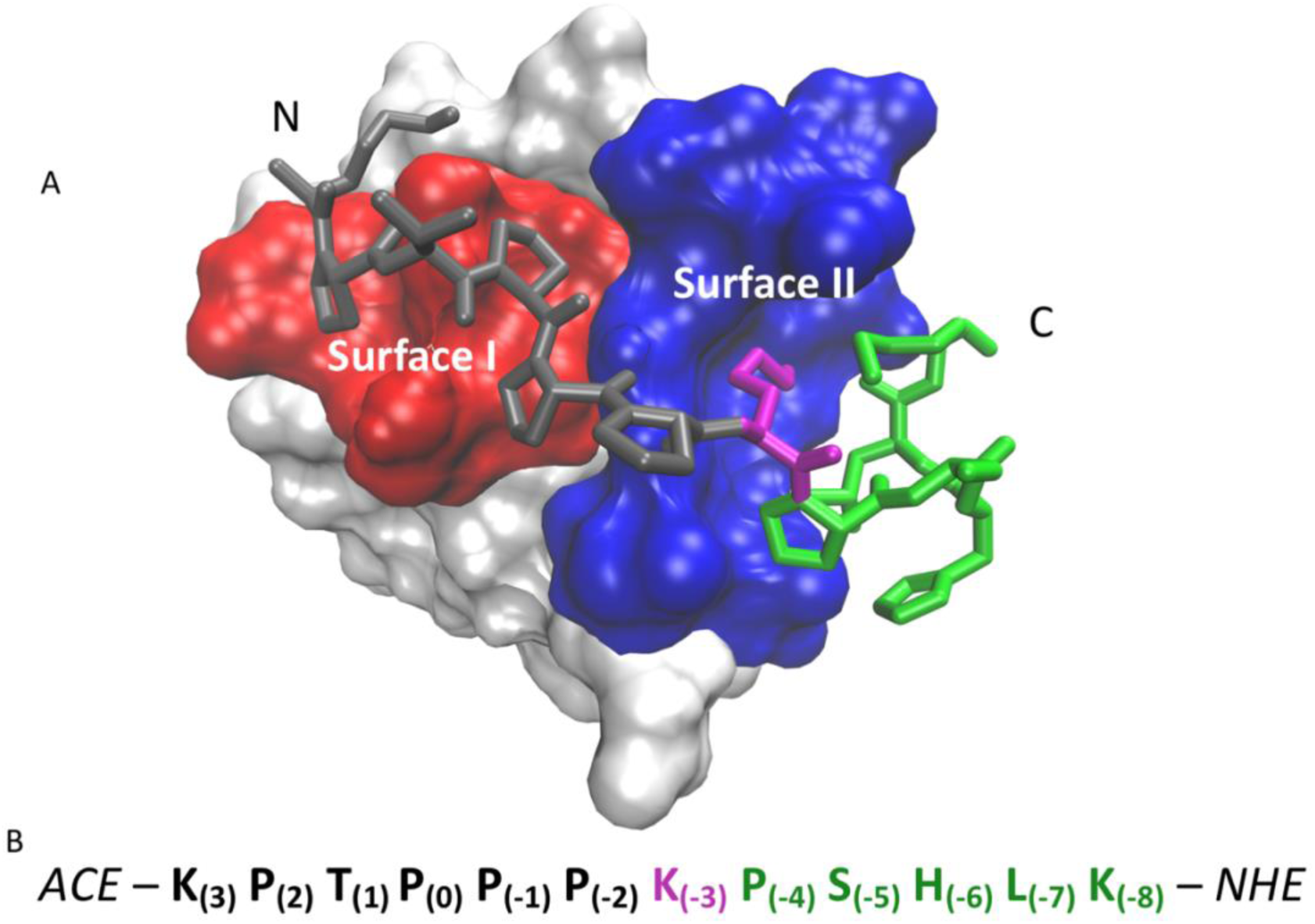
Description of system studied. A) Surface view of AbpSH3 bound to ArkA from NMR [72], showing the two binding surfaces (SI (red) and SII (blue)) with bound ArkA in gray (seg1), magenta (Lys(−3)) and green (seg2). The NMR structure was determined with a longer 17-residue ArkA sequence (residues 6 through −10) [73], but only the shorter ArkA sequence is displayed. The N and C-termini are labeled. B) Sequence of ArkA used in all simulations with seg1 shown in black, the central lysine in magenta, and seg2 in green. The capping groups on the C and N-terminal ends are also shown.

AbpSH3 has the typical SH3 fold with a five-stranded β-sandwich and long RT-loop, which is involved in ArkA binding [72]. The 12-residue ArkA peptide contains three Lys residues (Fig 1B), giving it a net positive charge, while the AbpSH3 domain has a net negative charge of −12. Thus, electrostatic attraction contributes to the affinity between the peptide and domain. ArkA can be described as two segments (seg1 and seg2) where seg1 is the N-terminal proline-rich end and seg2 is the C-terminal segment (Fig 1B). The proline-rich seg1 interacts with AbpSH3 in the typical class II orientation, with a PxxPx+ sequence that forms a PPII helix with each Px well packed into a groove [72]. The region of AbpSH3 which binds to the PxxP motif is referred to as surface I (SI) (Fig 1A). The C-terminal seg2 forms a 3^10^ helix in the NMR structure and makes contacts on a region of AbpSH3 distinct from SI, referred to as surface II (SII) (Fig 1A). The conserved Lys(−3) serves as the dividing residue between seg1 and seg2, and binds between SI and SII in a negatively charged ‘specificity pocket’, packing against a Trp side chain [72]. Previous NMR experiments have shown that seg1, containing the PxxP motif, can bind to AbpSH3 without seg2, but it does not fully engage the binding surface [40]. Seg2 alone, on the other hand, shows no detectable binding by NMR titration [40]. The role of each segment in the full binding pathway has not previously been investigated.

Using MD simulations, we found that ArkA initially forms a heterogeneous encounter ensemble, followed by the tight binding of seg1 and seg2 in the correct orientation with the formation of specific contacts (Fig 2). Significantly, ArkA forms many nonnative contacts in this encounter ensemble, but they are restricted to the canonical highly acidic binding surface of the AbpSH3 domain. Seg1 is largely pre-structured in a PPII helix and only needs to lock into the grooves of SI to bind, while seg2 is more conformationally flexible. The PPII helix in seg1 can bind in a reverse orientation in the encounter complex, and seg2 may be important for stabilizing the correct orientation of ArkA on the binding surface. Nonspecific hydrophobic and long-range electrostatic interactions stabilize the encounter complex, while specific hydrophobic interactions form only on transition from the encounter complex to the fully engaged state. Our simulations show that step 1 of binding is more than an order of magnitude faster than the overall association rate determined by NMR relaxation dispersion experiments. Our binding model also explains the greater influence of hydrophobic interactions on binding compared to long-range electrostatic interactions, which likely only affect step 1 of the binding pathway. Overall, we have gained an understanding of the different interactions of the two ArkA peptide segments with AbpSH3 along the binding pathway. This provides insights for how binding of this common interaction domain in other proteins may be tailored to meet their specific functional needs.

**Fig 2.**
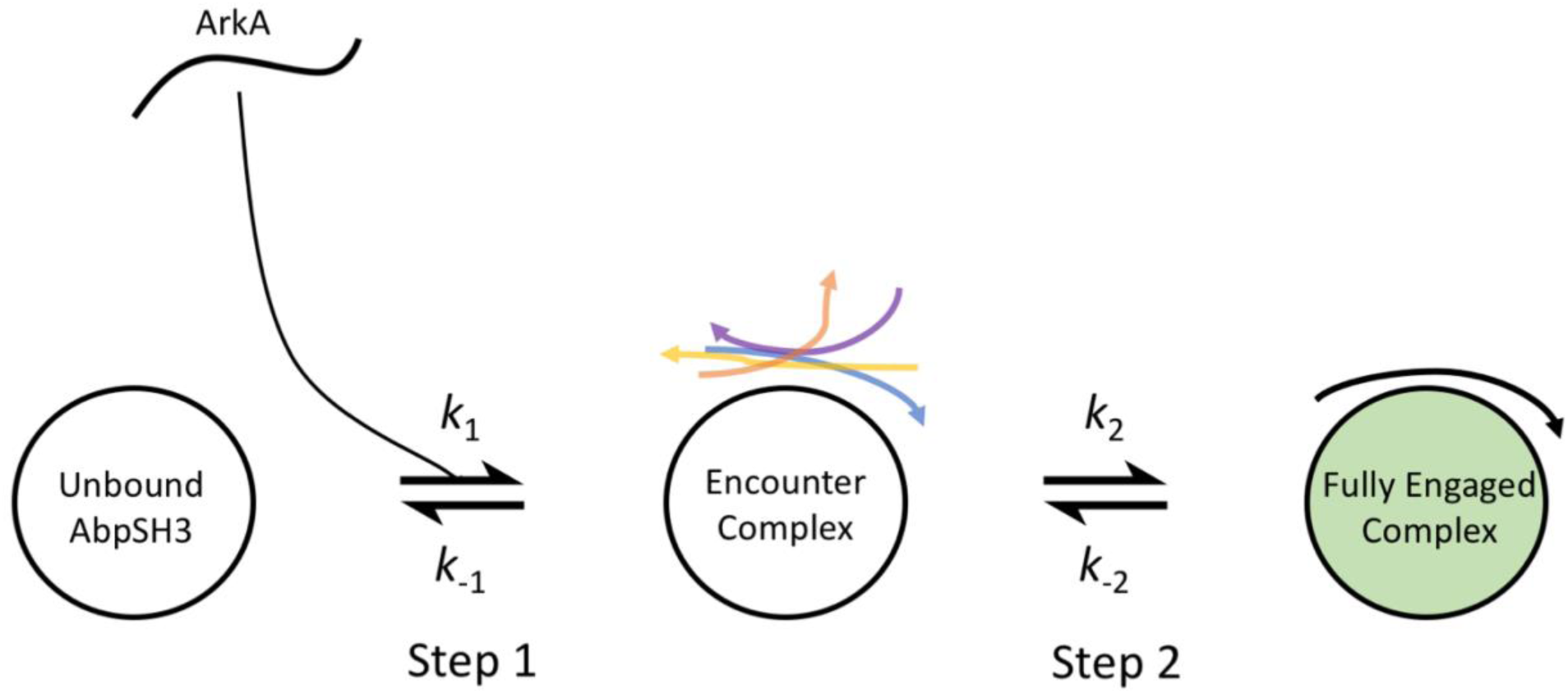
Proposed ArkA-AbpSH3 binding model. Two-step binding pathway with ArkA shown as a line and AbpSH3 as a circle. The heterogeneous encounter complex is proposed as an intermediate state between the unbound state and fully engaged complex. The forward and reverse rate constants for each step are labeled.

## Methods

### MD Simulations

MD simulations were run on four constructs: ArkA bound to AbpSH3 (bound simulations), ArkA alone (unbound simulations), ArkA binding to AbpSH3 (ArkA binding simulations), and ArkA seg1 binding to AbpSH3 (seg1 binding simulations). The bound simulations were all started from the lowest energy NMR structure of ArkA bound to AbpSH3 (PDB: 2RPN) [72]. Before running the simulations, the ArkA sequence was truncated to the 12-residue (Fig 1B) construct and a capping acetyl group was added to the ArkA N-terminus. Two different starting structures were used to initiate the unbound simulations. One starting structure was from the NMR structure of ArkA bound to AbpSH3 and the other was a fully extended peptide. For the ArkA and seg1 binding simulations, the peptide construct was placed at least 10 Å from AbpSH3 to ensure that the peptide and domain were not interacting at the beginning of the binding simulations (simulations were run with a non-bonded cutoff distance of 9 Å for the direct space sum). For the binding simulations, the starting structure of both ArkA and seg1 came from the ArkA unbound simulations, and the AbpSH3 structure came from the unbound crystal structure (PDB: 1JO8) [70]. The effective concentration of the protein in our simulations was around 4 mM (S2 Table), which is close to the experimental concentration of 1 mM. In all constructs, ArkA or seg1 were edited to have a capping acetyl group on the N-terminus (ACE) and a capping amide group on the C-terminus (NHE); except the bound simulations, which only have the acetyl group.

All simulations were run on Amber 16 using the Amber ff14SB forcefield [74], and the binding simulations were run with dihedral angle modifications that improve accuracy for the energy barrier between *cis* and *trans* states of the peptide bond [75]. The CUDA version of pmemd in Amber 16 was used to run the simulations on GPUs [76]. All simulations were solvated with TIP3P-FB water [77]. The bound simulations were solvated such that the edge of the box was at least 9 Å from any peptide or protein atom. Binding simulations were solvated with the edge at least 12 Å from any peptide or protein atom and adjusted to have an equal volume. The unbound simulations were solvated with water 15 Å from the edge of the peptide. The box dimensions are summarized in S3 Table. Salt ions were added to neutralize each system: 10 sodium ions for the seg1 binding and bound simulations, 9 sodium ions for the ArkA binding simulations and 3 chloride ions for the unbound simulations.

All systems were subject to two rounds of energy minimization of 1000 steps, where the first 500 steps were steepest descent and the second 500 steps conjugate gradient. The systems were then subject to heating from 100 to 300 K (40 ps with harmonic restraints with a force constant of 10 kcal/mol), and equilibration (50 ps with harmonic restraints with a force constant of 10 kcal/mol). All constructs, except the unbound simulations, were equilibrated again for 200 ps without restraints. Independent simulations were started with new random velocities. Bonds to hydrogen were constrained using the SHAKE algorithm during all simulations. The particle-mesh Ewald procedure was used to handle long-range electrostatic interactions with a non-bonded cutoff of 9 Å for the direct space sum.

The unbound simulations were run using temperature Replica Exchange MD [78]. 48 replicas were simulated using the NVT ensemble with temperatures from 290.00 - 425.00 K with geometric spacing to achieve similar exchange probabilities for all replicas (S1 Table) [78]. Each replica was equilibrated without restraints for 500 ps. The simulations were run with an integration step every 2 fs and coordinates stored every 25 ps. Three independent simulations from both an extended peptide and the conformation in the NMR structure were run for at least 125 ns each, resulting in a total of 1.15 μs of simulation at 300 K. The first 50 ns of each simulation was removed before analysis, resulting in 0.850 μs of simulation data used in the ArkA unbound ensemble.

The bound and binding simulations were run using the NPT ensemble at 300 K with a Monte Carlo barostat, new system volumes attempted every 100 steps, an integration step every 2 fs, and coordinates stored every 10 ps. The number and length of all simulations are summarized below (Table 1). The replica exchange simulations were run on the XSEDE resource Xstream [79], as well as a local cluster, and all other simulations were run on a local cluster.

**Table 1.** Summary of simulations run.

| **Construct** | **# of simulations** | **Length analyzed per simulation (ns)** | **Total length analyzed (μs)** |
| --- | --- | --- | --- |
| Unbound ArkA | 2 | 100 | 0.849 |
|  | 1 | 75 |  |
|  | 1 | 200 |  |
|  | 1 | 249.3 |  |
|  | 1 | 125 |  |
| Bound ArkA | 5 | 1832 | 9.16 |
| Binding ArkA | 50 | 1000 | 50 |
| Binding seg1 | 50 | 500 | 25 |

### Simulation Analysis

To analyze the trajectories, the AmberTools 16 package was used to measure dihedral angles, distances, secondary structure, solvent accessible surface area, hydrogen bonds, and salt bridges [76]. In house Python scripts were used for additional analysis. Error bars and standard deviations were calculated by computing independent values from each independent simulation and taking the standard deviation of those values.

#### Sampling Completeness

The running average of secondary structure per residue was used as a measure of completeness of sampling for the unbound simulations (S2 Fig). The autocorrelation time between replicas was also calculated to ensure the replicas were exchanging as expected (S3 Fig) [80].

#### Structural analysis of ensemble

Dihedral RMSD of ArkA from the NMR structure was calculated as described by Kreiger *et al*. for backbone dihedral angles [81]. The distance between the binding surface of AbpSH3 and ArkA was defined by the average of seven pairwise ArkA-AbpSH3 contact distances between *α* carbons where the SH3 domain residue changes chemical shift upon ArkA binding (S1 Fig) [40]. ArkA seg1 lacks three of these pairs, so the remaining four were used, in both cases this is called the binding surface distance. The dihedral angles were used to calculate the polyproline II helix (PPII) content as described by Masiaux *et al*. [82]. Residue distances were calculated based on the center of mass for each residue in ArkA and AbpSH3, and 8 Å was used as the cutoff distance to define a contact. Contact maps were created based on the percentage of the simulation during which residue contacts were made. Contact maps that describe a subset of the simulated ensemble (encounter, forward, reverse, seg2 only, encounter other, or unbound) were created based on the percentage of that subset that is making a contact.

The data from the binding simulations were divided into unbound, encounter complex, and fully engaged based on the binding surface distance (Table 2). The definition of the fully engaged complex was based on a binding surface distance less than 11.5 Å because the bound simulations had a binding surface distance less than 11.5 Å in 98% of the simulated ensemble. There is a clear free energy barrier between the fully engaged state and the encounter complex based on a population histogram from our binding simulations (S17 Fig). However, in our simulations, there appears to be no free energy barrier between the unbound state and the encounter complex, indicating that formation of the initial encounter complex is downhill in free energy. We still wanted to define the encounter complex separate from the unbound state in order to characterize this intermediate state to binding, so we chose the encounter complex definition to be between 11.5 and 23 Å binding surface distance. This definition of the encounter complex captures the states that are most populated along the binding surface distance reaction coordinate (S17 Fig), and also excludes states where ArkA has no contacts with the SH3 domain (S18 Fig), which exist at binding surface distances greater than 23 Å, defined as unbound. Within the encounter complex ensemble, four categories were defined based on contacts between ArkA and AbpSH3 (Table 2).

**Table 2.** Categories in binding simulations.

| **Binding Interaction** | **Contacts** |
| --- | --- |
| Unbound | Binding surface distance > 23 Å |
| Fully engaged | Binding surface distance < 11.5 Å |
| Encounter Complex | 11.5 Å < binding surface distance < 23 Å |
| Encounter forward | K(3) to AbpSH3 8, 9, or 10 and K(-3) to AbpSH3 33, 35, or 36 |
| Encounter reverse | K(-3) to AbpSH3 8, 9, or 10 and K(3) to AbpSH3 33, 35, or 36 |
| Encounter seg2 only | K(-3) to AbpSH3 33, 35, or 36<br>and K(-8) to AbpSH3 14, 15,16,17 or 49 or L(-7) to AbpSH3 32, 33, 36, 49<br>and not forward or reverse |
| Encounter other | All other structures |

The ArkA simulated ensembles were divided into natively folded conformations and nonnative conformations based on a histogram of the ArkA dihedral RMSD from the unbound simulations. The first minimum in the histogram, at 33.7° was used as a cutoff (S10 Fig). All ArkA conformations with a dihedral RMSD less than 33.7° are considered to have a native conformation, and those with a dihedral RMSD greater than 33.7° are considered to have a nonnative conformation. For the seg1 dihedral RMSD, a cutoff of 38.1° was used to define the native and nonnative conformations (S9 Fig).

Long-range electrostatic interactions between ArkA and AbpSH3s were analyzed by calculating the distance between the charged groups of the positively charged residues (N_ζ_ of Lys) on ArkA and the negatively charged residues (C_γ_ of Asp and C_δ_ of Glu) on AbpSH3. The calculation was performed on the ArkA binding simulations and bound simulations. A 10-Å cutoff distance was used to define an electrostatic interaction. An in-house python script was used to calculate the percentage of time each electrostatic interaction was present and the average number of electrostatic interactions present simultaneously for each simulation. An interaction had to be present in one of the simulations for at least 10% of the ensemble to be included in the results.

Hydrophobic contacts were selected from those hydrocarbon groups that are closest together in the NMR structural ensemble (2RPN) [72]. Contacts were defined based on a 6 Å cutoff distance between hydrocarbon groups. Hydrogen bonds were counted when the distance between the acceptor atom and donor heavy atom was less than 3 Å and the angle between the acceptor atom, donor hydrogen, and donor heavy atom was greater than 135°. Similarly, salt bridges were counted when the distance between the heavy atoms of the charged groups was less than 3 Å and the angle between the oxygen atom, hydrogen, and nitrogen was greater than 135°.

#### SH3 domain dipole moment

The net dipole of the AbpSH3 domain was calculated using the Protein Dipole Moments Server administered by the Weizmann Institute Department of Structural Biology [83] by uploading the crystal structure of the domain (PDB: 1JO8) [70].

#### Calculation of *k*_on_ and *k*_1_

The time constant, *τ*, for binding of ArkA and seg1 to AbpSH3 is related to the rate constant, *k*_on_ or *k*_1_, and the concentration of protein and peptide by the equation

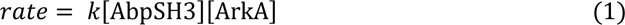

where *k* is either *k*_on_ or *k*_1_. *k*_on_ is the rate constant for complete binding to the fully engaged complex, while *k*_1_ is the rate constant for formation of the encounter complex. The volume of the boxes varied slightly between the two ArkA constructs, so the concentrations and rates were slightly different (S2 Table).

If the AbpSH3 concentration is held constant, then the transition from unbound to either the encounter complex or the fully engaged complex can be treated as a first-order reaction, so the binding time follows a Poisson distribution [84, 85]. The binding time constant, *τ*, was calculated from a fit of the empirical cumulative distribution function to the theoretical cumulative distribution function (TCDF) [85],

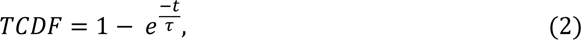

where *t* is time of the simulation when binding occurs. Since we observed some overlap in the distribution of binding surface distances for fully engaged and encounter complexes, we used a more stringent definition of binding for identifying transitions between states. To go from the encounter complex to the fully engaged complex we required that the binding surface distance be below 10.5 Å for at least 1 ns, and to go from unbound to encounter we required it to be below 21 Å for 1 ns. *τ* was then used to calculate the binding rate constant,

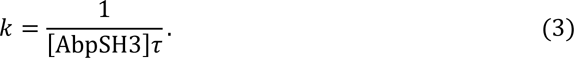

The standard deviation in calculation of *k*_1_ was determined using the bootstrap method, but we could not determine a standard deviation for *k*_on_ because not all simulations reached the fully engaged state.

Our simulations were performed without salt present (aside from neutralizing ions), while the experimental rate constants were measured with 100 mM NaCl and 50 mM phosphate, which could affect the binding rate, particularly for the formation of the encounter complex, which is partially driven by electrostatic interactions.

To determine the number of transitions from the encounter complex, we used a similar method and required that the binding surface distance be above 25 Å for at least 1 ns for the transition from encounter complex to unbound to be counted in order to make sure that we only counted true transitions out of the encounter complex free energy minimum (S17 Fig).

### Experimental

The AbpSH3 protein and ArkA peptide were produced as previously [40]. A set of _15_N Carr-Purcell-Meiboom-Gill (CPMG) relaxation dispersion experiments were recorded on 1 mM _15_N labeled domain with ∼9% bound to unlabeled peptide in 50 mM phosphate, 100 mM NaCl, pH 7.0 at 10 °C. The data were collected at two different static magnetic field strengths (500 and 800 MHz) generating a series of 2 x 21 2D _1_H-_15_N correlation maps measured as a function of CPMG frequency (S20 Fig). The spectra were processed using standard approaches and the program chemex [86, 87] assuming a 2-state reaction [40, 88], which has been applied to a few other domains [31, 38]. Global fitted values of *k*_ex_ and *p*_bound_ were extracted (224 s^-1^ and 0.08) from these data. A value of *k*_off_ was subsequently calculated (206.5 s^-1^) using the equation

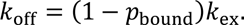

A value of *k*_on_ (1.21 x10^8^ M^-1^ s^-1^) was then calculated from the K_d_ value of 1.7 μM [40] using the expression *k*_off_/K_d_.

## Results & Discussion

### ArkA disorder when unbound & bound

We first sought to characterize the structural ensemble of the unbound ArkA peptide to help determine how the intrinsic structural propensities of ArkA contribute to binding. In order to determine the structural ensemble of the unbound ArkA peptide, we simulated the 12-residue ArkA alone using REMD [78] (unbound simulations). The completeness of sampling was examined using the running averages of 3^10^ helix, bend, and turn structure, as well as end-to-end distance (S2 Fig). The efficiency of exchange between the replicas was confirmed by determining that the autocorrelation time for the replica temperature (time constant 7-10 ns) was well below our simulation time (S3 Fig). The structural ensemble of ArkA simulated alone shows that it is behaving as a disordered peptide with both the end-to-end distance and dihedral angle RMSD sampling multiple states that are different from the NMR reference structure (Fig 3A).

**Fig 3.**
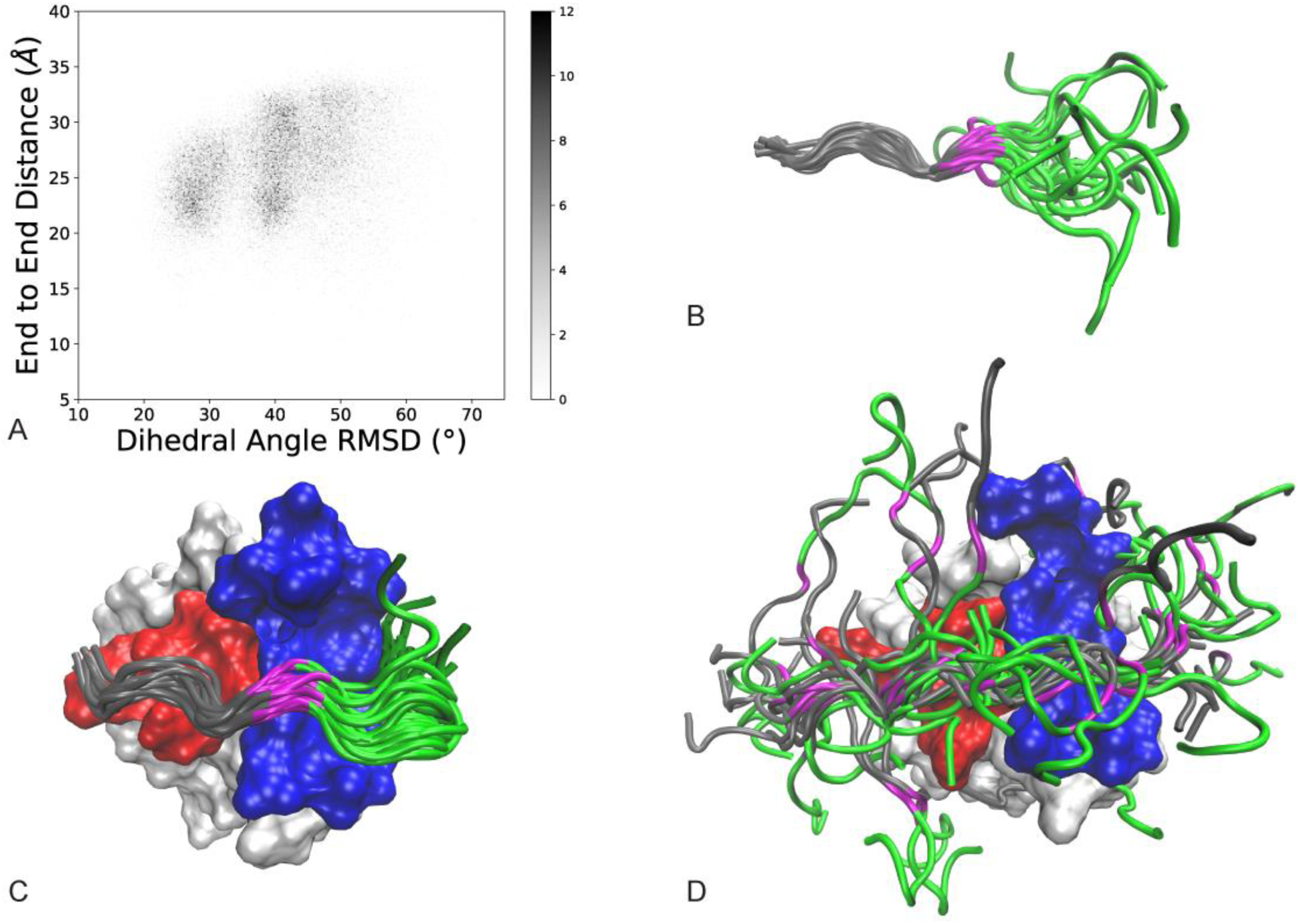
Characterization of unbound, bound, and encounter ensembles. A) Conformational ensemble of unbound ArkA from REMD simulations. End-to-end distance is the distance between the C and N-terminal ends of ArkA and dihedral angle RMSD is calculated for ArkA with the lowest energy NMR structure (2RPN) as the reference [72]. Darker shading indicates a larger fraction of the total ensemble, as indicated by the color bar. B) Overlay of 13 randomly selected ArkA conformations from unbound simulations with seg1 residues Pro(2) to Pro(−2) backbone aligned. C) Overlay of 15 randomly selected ArkA conformations from bound simulations with the SH3 domain aligned. D) Overlay of 38 randomly selected ArkA conformations in the encounter complex from ArkA binding simulations with the SH3 domain aligned. AbpSH3 SI is shown in red and SII in blue. ArkA is shown in in gray (seg1), magenta (Lys(−3)) and green (seg2).

We also ran simulations starting from the NMR bound structure of ArkA to compare its structure when fully engaged with AbpSH3 (bound simulations) (Fig 3C). NMR experiments have shown that in the fully engaged state ArkA seg1 adopts a PPII helix and seg2 is often in a 3^10^ helix (S5 Fig) [72]. In both the alone and bound simulations, seg1 of ArkA is largely structured with the majority of time spent in a PPII helix, while seg2 is less structured (Fig 4). Even in the bound simulations seg2 is only in a 3^10^ helix 35% of time, indicating that the majority of the diversity in ArkA secondary structure occurs in the seg2 region. Contact maps between ArkA and AbpSH3 also show that the entire peptide is more flexible in the bound simulations than the NMR structures indicate. Both segments of ArkA have fewer contacts with AbpSH3 in the simulated ensemble than the NMR ensemble, especially seg2 (S7 Fig).

**Fig 4.**
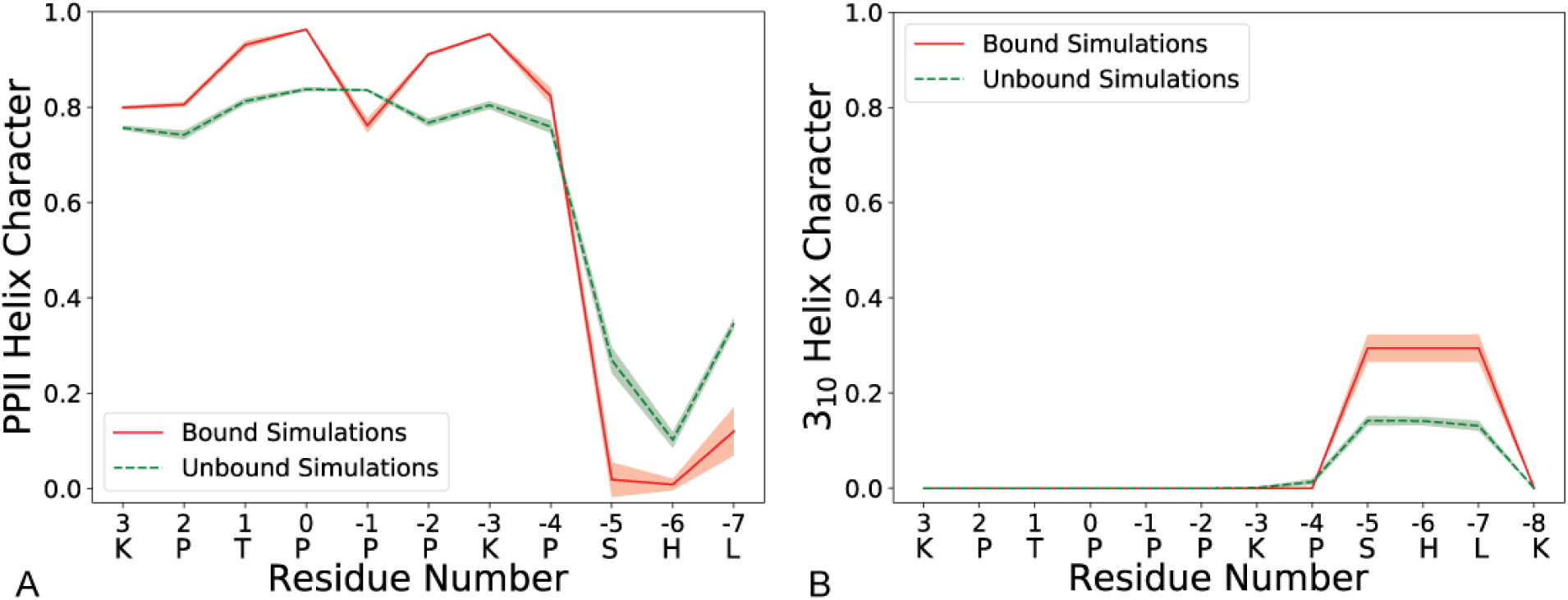
Quantifying bound and unbound secondary structure. Fraction of time each ArkA residue spends in PPII Helix (A) or 3^10^ Helix (B) during the bound and unbound simulations. The one letter codes for ArkA residues are included on the x-axis. The shaded region represents the standard deviation between independent simulations.

### ArkA binds via an encounter complex intermediate

After determining the conformational ensemble of the unbound ArkA peptide, we wanted to characterize the binding pathway by running ArkA binding simulations. To start the 1-μs ArkA binding simulations, we chose five structures with differing values of dihedral RMSD and end-to-end distance (S8 Fig) and used these to initiate 50 independent binding simulations (10 from each structure) as indicated in Table 1. This ensured that the binding simulations would not be biased by a single starting ArkA conformation. We also ran binding simulations with the shorter seg1 peptide (seg1 binding simulations), which contains the PxxP motif that is relatively structured, to examine the different roles of seg1 and seg2 in binding (S1 Text).

In the ArkA binding simulations, we found that an initial encounter complex forms quickly before ArkA transitions more slowly to a fully engaged state (S1 Movie and S2 Movie). As described in the methods, the fully engaged state was defined as a structure where the binding surface distance is below 11.5 Å, while we defined the encounter complex as having binding surface distance between 11.5 Å and 23 Å. In the binding simulations, ArkA passes through the encounter complex (Fig 5A-B) before reaching the fully engaged state. Interestingly, in some of the independent simulations ArkA forms an encounter complex, then dissociates before rebinding (Fig 5A), while in others it quickly reaches the same fully engaged state observed in the bound simulations (Fig 5B). In the seg1 binding simulations we similarly observed the formation of an initial encounter complex, followed by either unbinding and rebinding, or transition to a stable fully engaged bound state (S11 Fig), while in the bound simulations, the complex remains in the fully engaged state 98% of the time (Fig 5C).

**Fig 5.**
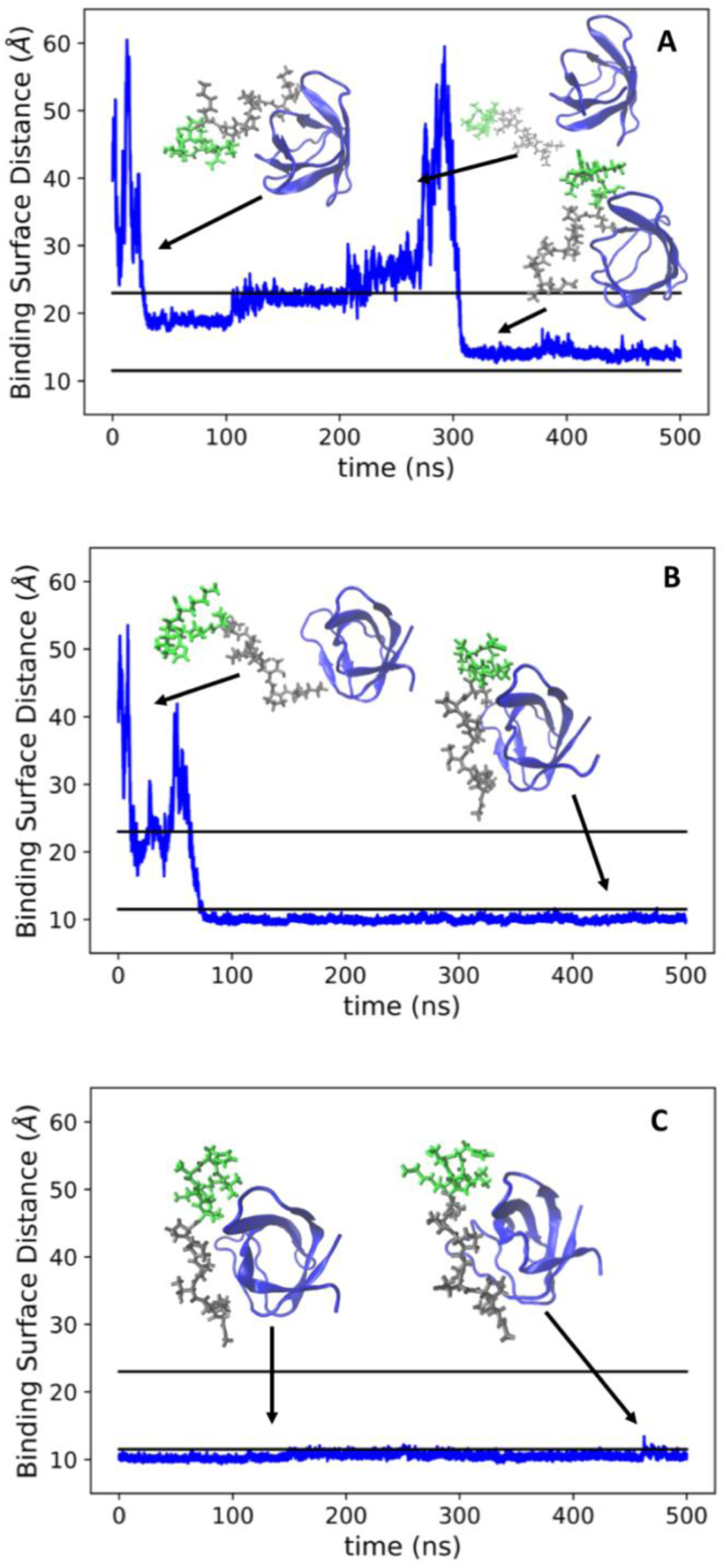
Time traces of binding and bound simulations. Distance between ArkA and the binding surface of AbpSH3 for the first half of two example ArkA binding simulations (A, B) and one bound simulation (C). The black lines correspond to our definition of the encounter complex (23 Å) and the fully engage complex (11.5 Å). In the ArkA binding simulations, the largest box dimension is 80 Å (S3 Table) and the maximum AbpSH3 domain diameter is 33 Å. Center of mass distances between ArkA and the SH3 domain range from 11 to 64 Å in the binding simulations, while binding surface distances range from 9 to 67 Å. In the bound simulations, the binding surface distances range from 8 to 16 Å.

### The ArkA encounter complex is a heterogeneous ensemble that includes nonnative interactions

To further examine the nature of the encounter complex, we projected the data onto coordinates corresponding to native backbone folding (dihedral RMSD) and binding (pairwise binding surface distance) (Fig 6). In the binding simulations, ArkA samples many states with different degrees of native folding and binding before reaching the fully engaged and native folded state found in the lower left of the plot (Fig 6A, blue rectangle). In particular, the encounter complex ensemble is a highly heterogeneous state, as shown in Fig 3D, and 57% of the ArkA binding ensemble occupies the encounter complex without forming the native ArkA fold (Fig. 6A), indicating that ArkA does not need to be already preformed in the native conformation before interacting with AbpSH3, consistent with an induced-fit binding mechanism. However, in 20% of the binding ensemble ArkA has a native fold but is still in the encounter complex. This indicates that, at times, ArkA may first adopt a native fold and then reorient and dock into the fully engaged state in a conformational selection mechanism. Multiple steps and potential pathways to binding may exist within the encounter complex ensemble. Additionally, Fig 6B confirms that the bound simulations stay fully engaged but do rarely (4% of the ensemble) sample nonnative conformations that are different from the NMR structure (unfolded and fully engaged in Fig 6B). We only observe two brief instances, totaling to less than 2% of the ensemble, where the complex transitions to the encounter complex and back to the fully engaged state in one of the five bound simulations.

**Fig 6.**
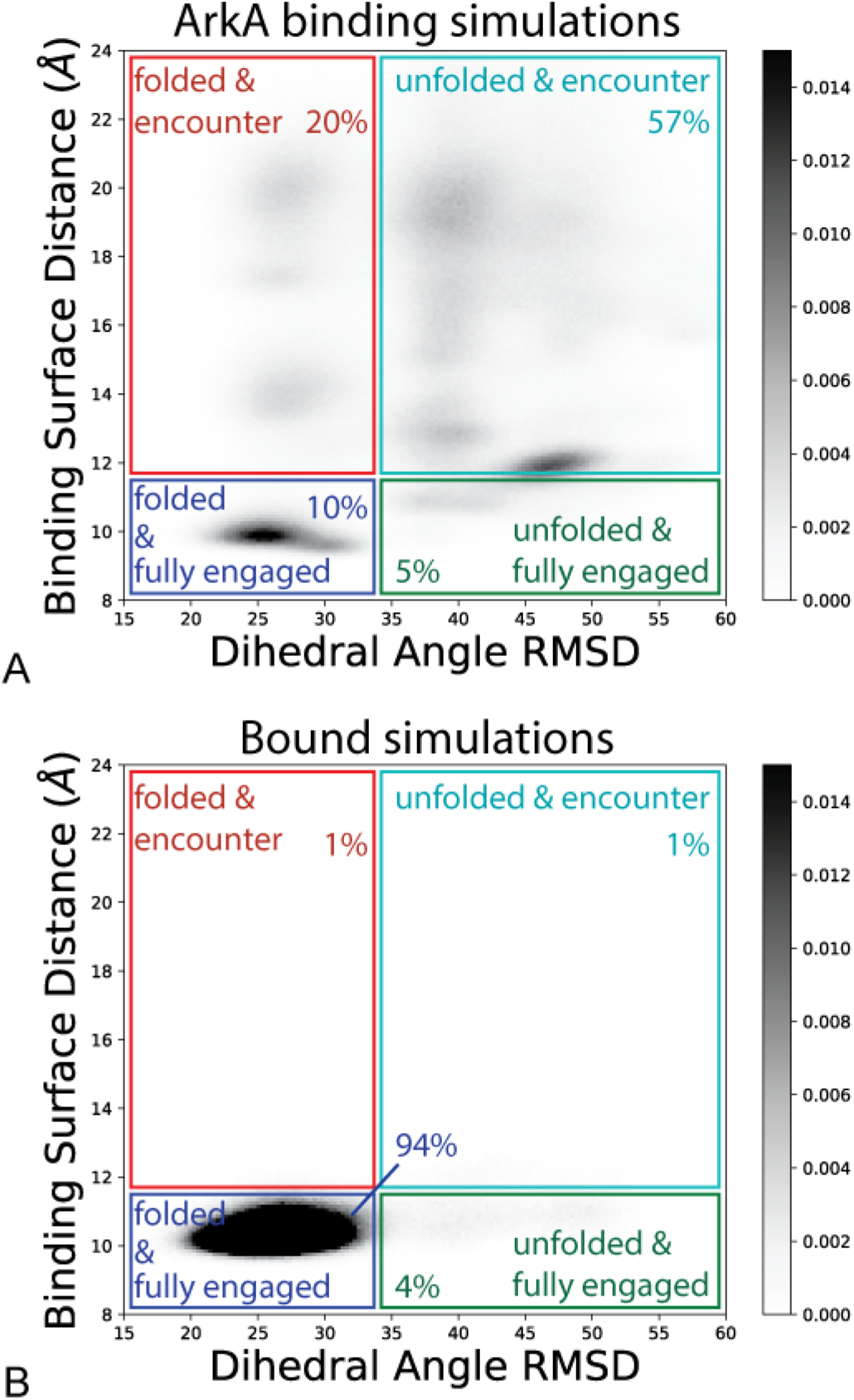
ArkA states sampled during binding and bound simulations. Distance between ArkA and the binding surface of AbpSH3 graphed against the ArkA backbone dihedral angle RMSD for ArkA binding simulations (A) and bound simulations (B). Darker shading indicates a larger fraction of the total ensemble, as indicated by the color bar. Colored boxes partition the ensemble into four states: folded and fully engaged (blue), unfolded and fully engaged (green), folded and encounter (red), unfolded and encounter (cyan). In this figure, folded refers to the native NMR structure fold (dihedral RMSD less than 33.7°), and unfolded refers to a nonnative conformation different from the NMR fold (dihedral RMSD greater than 33.7°). Percentages indicate the occupancy of each state in the overall simulated ensemble. In the ArkA binding simulations, 8% of the ensemble is in the unbound state, which is not shown on the plot.

Contact maps also illustrate the heterogeneous nature of the encounter complex ensemble (Fig 7). In the encounter ensemble, ArkA makes nonnative contacts which are not seen in the bound simulations. These contacts are mainly on the two binding surfaces (SI and SII), indicating that electrostatic orientational steering guides the positively charged ArkA to interact with the correct surface of the domain, though not necessarily with native contacts. This is explained by the presence of a net dipole moment on the SH3 domain of 242 Debyes, with the negative end of the dipole located at the binding surface and the positive end on the opposite side of the domain, as has been observed for other SH3 domains [89]. In the encounter complex, seg1 forms more contacts (9 on average) with the binding surface than seg2 (6 on average).

**Fig 7.**
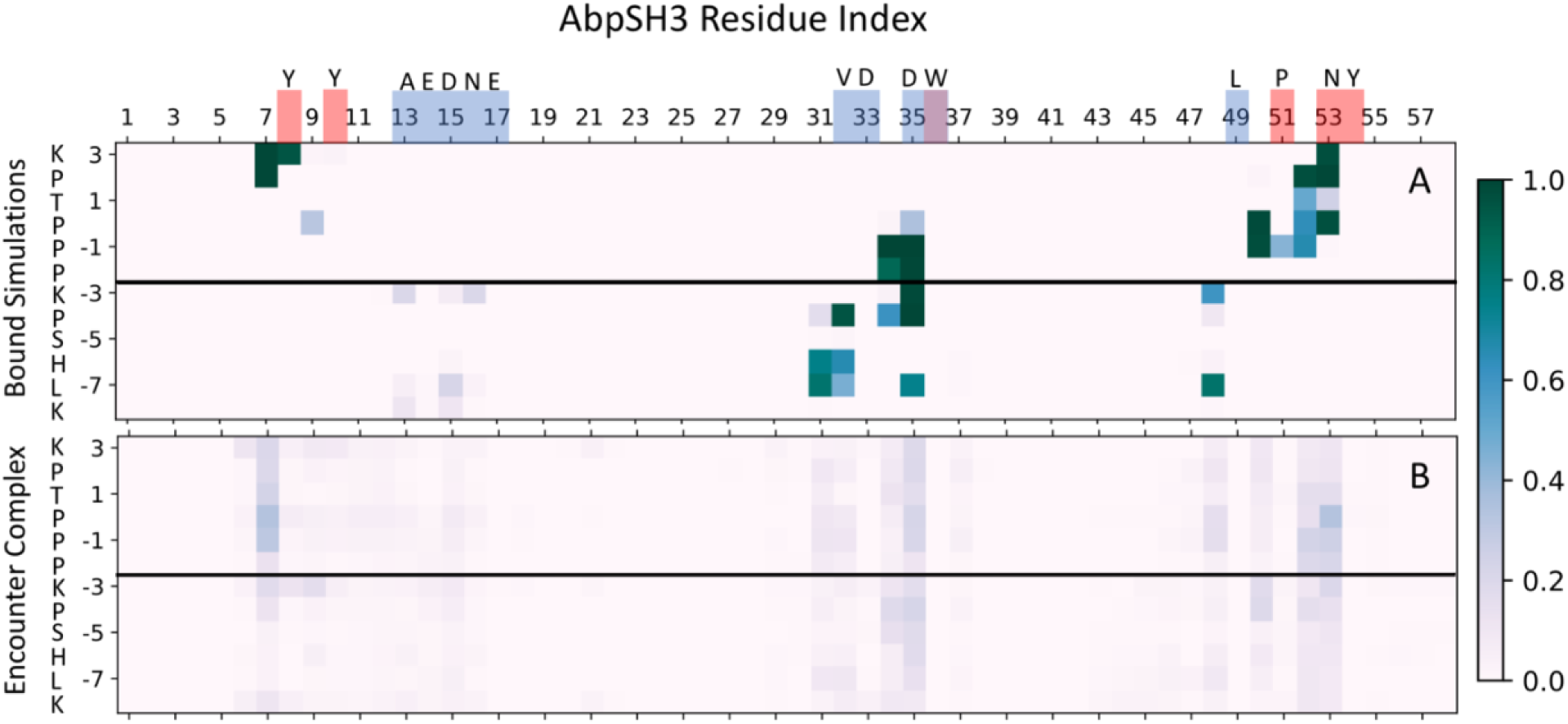
Contact maps between AbpSH3 and ArkA. Contact maps of the fully engaged state from the bound simulations (A) and the encounter complex ensemble from the binding simulations (B). The shade of the square indicates the fraction of the ensemble with that contact. The red and blue squares along the AbpSH3 residue index show which amino acids are in SI and SII, respectively. The black line indicates the separation of seg1 and seg2, and the single letter amino acid codes are included for ArkA and the residues in SI and SII. Several contacts are formed part of the time in the encounter complex but are not occupied at all in the bound simulations, indicating that nonnative contacts are part of the encounter complex ensemble. On average in the encounter complex, seg1 is in contact with 9 SH3 domain residues, while seg2 is in contact with 6 residues. ArkA K(−3) is in contact with 2 SH3 domain residues on average in the encounter complex, consistent with its role as an important central residue for binding.

The nonnative contacts on the binding surface that are formed in the encounter complex are consistent with ArkA binding in reverse in part of the encounter ensemble (Fig 8). In general, SH3 domains depend on the PxxP motif to bind, and as this is a pseudo-palindromic motif, reverse binding for seg1 on SI is not surprising. To further examine the conformational states in the encounter complex, we broke the encounter complex into four categories (defined in the methods, Table 2): forward, reverse, seg2 only, and other (Table 3). The reverse structures are found in the part of the encounter ensemble that has a binding surface distance higher than 15 Å (Fig 6A), while the forward structures are found at binding surface distances less than 15 Å, as expected. The encounter complex contact map (Fig 7) shows that seg1 interacts more with the binding surface overall than seg2. Furthermore, Table 3 shows that the encounter complex is twice as likely to sample a state with seg1 engaged in the correct orientation on SI than a state with only seg2 engaged in the correct place on SII. Together, this indicates that seg1 likely binds before seg2. Seg2 may be needed to ensure specific forward binding since seg2 does not interact with the domain when ArkA binds in reverse (Fig 8). Both the ArkA and seg1 binding simulations exhibit forward and reverse binding, showing that in the encounter complex the two segments behave somewhat independently. However, the ArkA encounter complex ensemble is much more complex and heterogeneous than that of the short seg1 peptide (S12 Fig), and ArkA samples the forward state less often than the seg1 peptide (Table 3), indicating that this short peptide may not give an accurate representation of how that segment behaves as part of the longer sequence.

**Fig 8.**
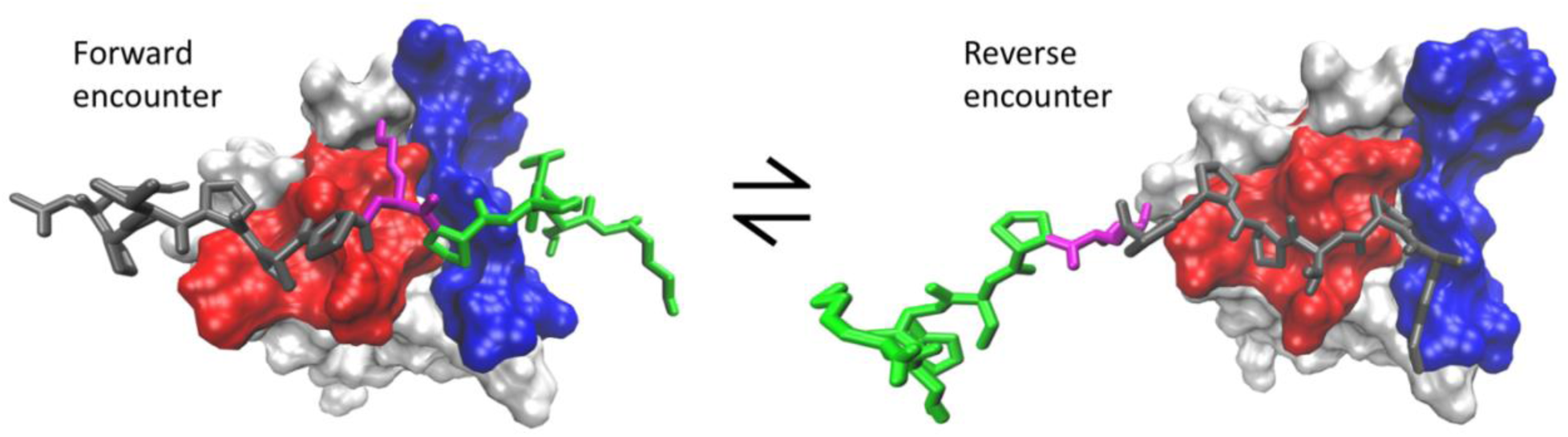
Forward and reverse representations of ArkA during binding. Snapshots from MD simulation showing both the forward and reverse orientations of ArkA that are possible during binding. ArkA is shown in the stick representation with seg1 in grey, seg2 in green, and K(−3) in magenta. AbpSH3 SI is shown in red and SII in blue. The double headed arrow signifies that the ArkA orientation can flip during a simulation.

**Table 3.**
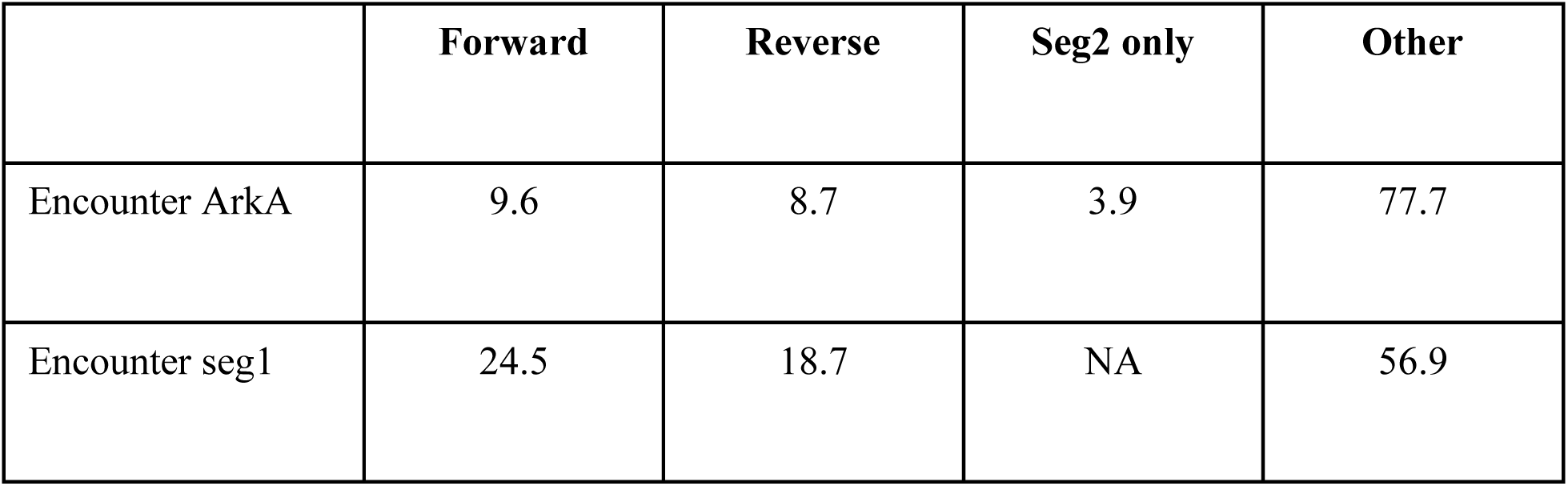
Percentage of the encounter complex ensemble in each category for ArkA and seg1 simulations.

Although the percent of the encounter complex ensemble that is in the forward and reverse encounter is about the same (Table 3), 33 of the 50 individual binding simulations sampled forward encounter at some point in the simulation, compared to only 15 that sampled the reverse encounter. We found that generally, when the encounter complex enters the forward encounter state it does not change orientation; however, sometimes ArkA spins around and shifts over to go from forward to reverse (2 times out of 31) or reverse to forward (4 times out of 15) without entering the unbound state in between (S3 Movie). The encounter complex is in dynamic exchange between different predominantly nonnative conformations and contacts, including the forward and reverse orientation of the peptide (Fig 3D). This dynamic exchange may help to prevent the encounter complex from being trapped in off-pathway states for binding, such as a reverse encounter complex.

The nonspecific, disordered encounter complex that we characterize from our simulations is similar to encounter complexes seen previously in MD simulation studies of the proline-rich Sos peptide binding to the c-Crk N-SH3 domain [21,23,56]. Ahmad *et al.* performed very short binding simulations where the Sos-SH3 domain complex formed very rapidly, and they identified three binding modes, including forward and reverse orientations of the PPII helix on the binding surface [21]. Because these binding modes formed very quickly, they likely represent different conformations within the encounter complex ensemble, rather than fully engaged states of the complex. In our simulations we have been able to sample the encounter complex ensemble more extensively and quantify the occupancy of different conformational states within this ensemble. Ahmad *et al.* also identified a binding mode where the Sos peptide interacts with the c-Crk N-SH3 domain at a new binding surface [21]; however, our simulations do not show any evidence that the ArkA peptide interacts significantly with a surface of the AbpSH3 domain other than the canonical binding surface. The nonnative interactions of ArkA with the AbpSH3 binding surface we observe in the encounter complex ensemble are similar to the alternate states observed by Yuwen *et al.* in simulations characterizing a mutant c-Crk N-SH3 domain interacting with the Sos peptide (designed to imitate the encounter complex) [56]. With our comprehensive analysis, it now seems likely that a diverse encounter complex ensemble that includes nonnative interactions may be characteristic of the binding between proline-rich peptides and SH3 domains.

### Long-range electrostatic interactions stabilize the encounter complex

Because of the complementary charges of ArkA and AbpSH3 and previous studies that focused on electrostatic interactions, we chose to particularly examine the role that long-range electrostatic interactions play in the encounter complex. We measured the intermolecular electrostatic contacts present in the encounter complex ensemble and in the fully engaged ArkA-AbpSH3 complex. While the long-range electrostatic contacts present in the encounter complex ensemble are more diverse (nonspecific) than those in the bound simulations (Fig 9), the average number of electrostatic contacts in the encounter complex ensemble at any given time is very similar to the average number in the bound simulations (Table 4). Thus, the main favorable contribution of the positively charged ArkA peptide interacting with the negatively charged AbpSH3 binding surface is gained upon formation of the encounter complex rather than upon transitioning from the encounter to the fully engaged state.

**Fig 9.**
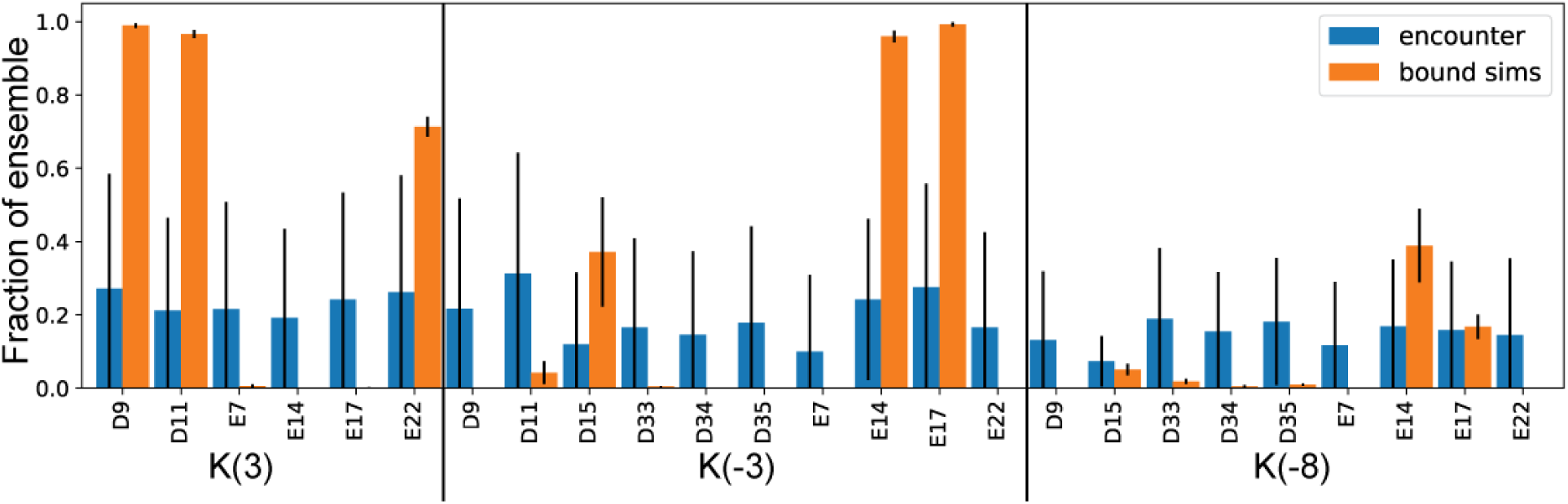
Long-range electrostatic interactions are non-specific in the encounter complex and specific in the bound simulations. Frequency of particular long-range electrostatic interactions in the ArkA encounter complex (blue bars on the left) and ArkA bound simulations (orange bars on the right). The large labels indicate the ArkA residue involved in the long-range electrostatic interaction and the small labels indicate the AbpSH3 residue. Error bars represent the standard deviation between independent simulations.

**Table 4.**
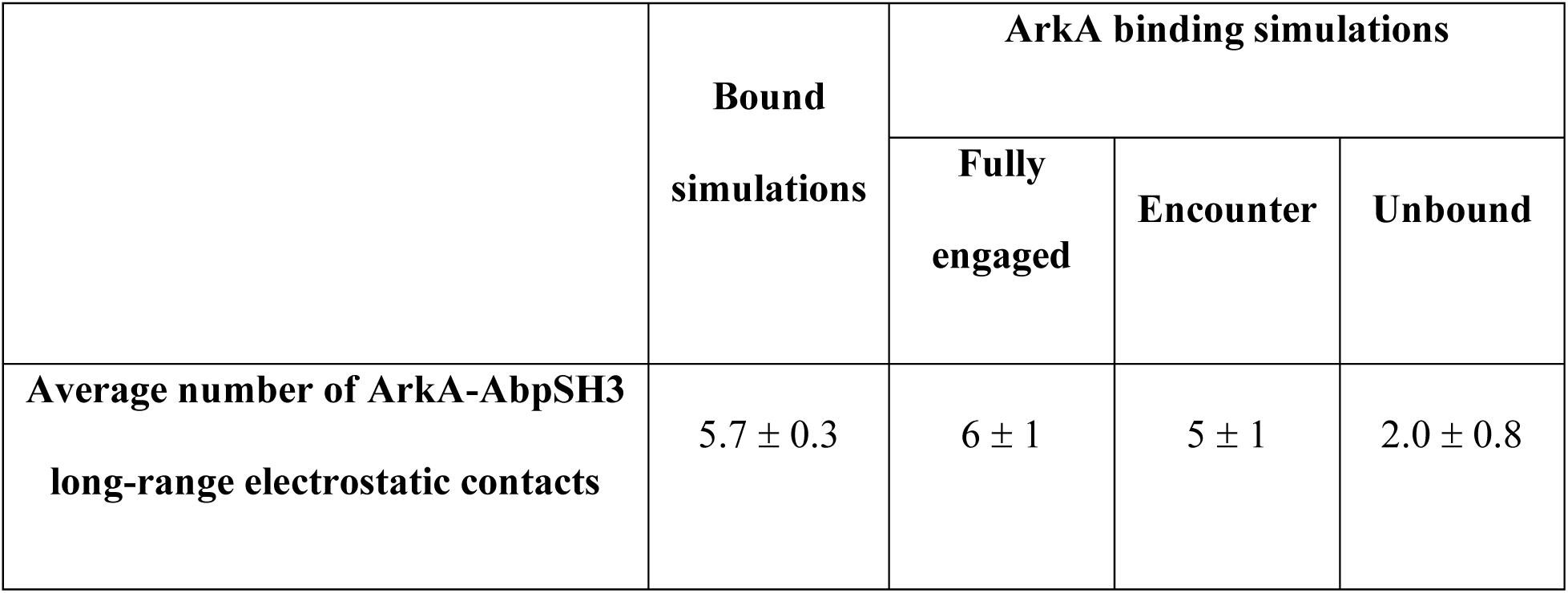
Average number of long-range electrostatic contacts for different states.

|  | **Bound**<br>**simulations** | **ArkA binding simulations** |  |  |
| --- | --- | --- | --- | --- |
|  |  | **Fully**<br>**engaged** | **Encounter** | **Unbound** |
| **Average number of ArkA-AbpSH3**<br>**long-range electrostatic contacts** | $5.7 \pm 0.3$ | $6 \pm 1$ | $5 \pm 1$ | $2.0 \pm 0.8$ |

Previous studies of SH3 binding have also found that electrostatic interactions are important for the formation of the complex [89]. MD simulations of the Sos peptide binding to c-Crk N-SH3 were able to specifically identify electrostatic interactions that occur in the encounter complex, including nonnative contacts, although they did not quantify these interactions [21, 23]. Experimental studies of the viral NS1 peptide binding to the CrkII N-SH3 domain indicate that electrostatic contacts are important for specific binding, and that flexibility in the fully engaged state allows increased electrostatic stabilization as multiple interactions form as part of the ensemble of bound states [68, 89]. Our simulations indicate that a diversity of different electrostatic contacts, each present in only part of the ensemble, is even more characteristic of the ArkA-AbpSH3 encounter complex than the fully engaged complex. The heterogeneity, or ‘fuzziness’, of the encounter complex ensemble is important, as there can be multiple pathways from this fuzzy encounter state to the fully engaged complex [37]. Electrostatic interactions can enhance this effect, not only stabilizing the encounter complex, but also lowering the free energy barrier to transition between basins and transition to the fully engaged state [13, 37]. As many of the ArkA-AbpSH3 encounter complex electrostatic contacts do not form at all in the fully engaged ensemble, it is also important that none are strong enough interactions to trap the complex in a conformation incompatible with transitioning to the fully engaged state.

### Hydrophobic and short-range interactions are nonspecific in the encounter and specific in the fully engaged complex

Since simulations indicate that long-range electrostatic interactions are already formed in the ArkA-AbpSH3 encounter complex, we sought to identify what energetically favorable changes occur upon transitioning from the encounter complex to the fully engaged state. By measuring the solvent accessible surface area of the complex, we found that in the encounter complex part of the SH3 domain binding surface is buried (Fig 10A) because ArkA forms transient nonspecific hydrophobic interactions with the binding surface. However, in transitioning to the fully engaged complex, the ArkA PPII helix packs into the grooves in SI, and native contacts that are largely absent in the encounter ensemble form between hydrophobic sidechains at the interface (Fig 10C). This buries more of the binding surface, which transitions from ∼50 to ∼45 to ∼40 nm^2^ solvent exposed surface area as the complex transitions from unbound to the encounter complex to the fully engaged complex (Fig 10A).

**Fig 10.**
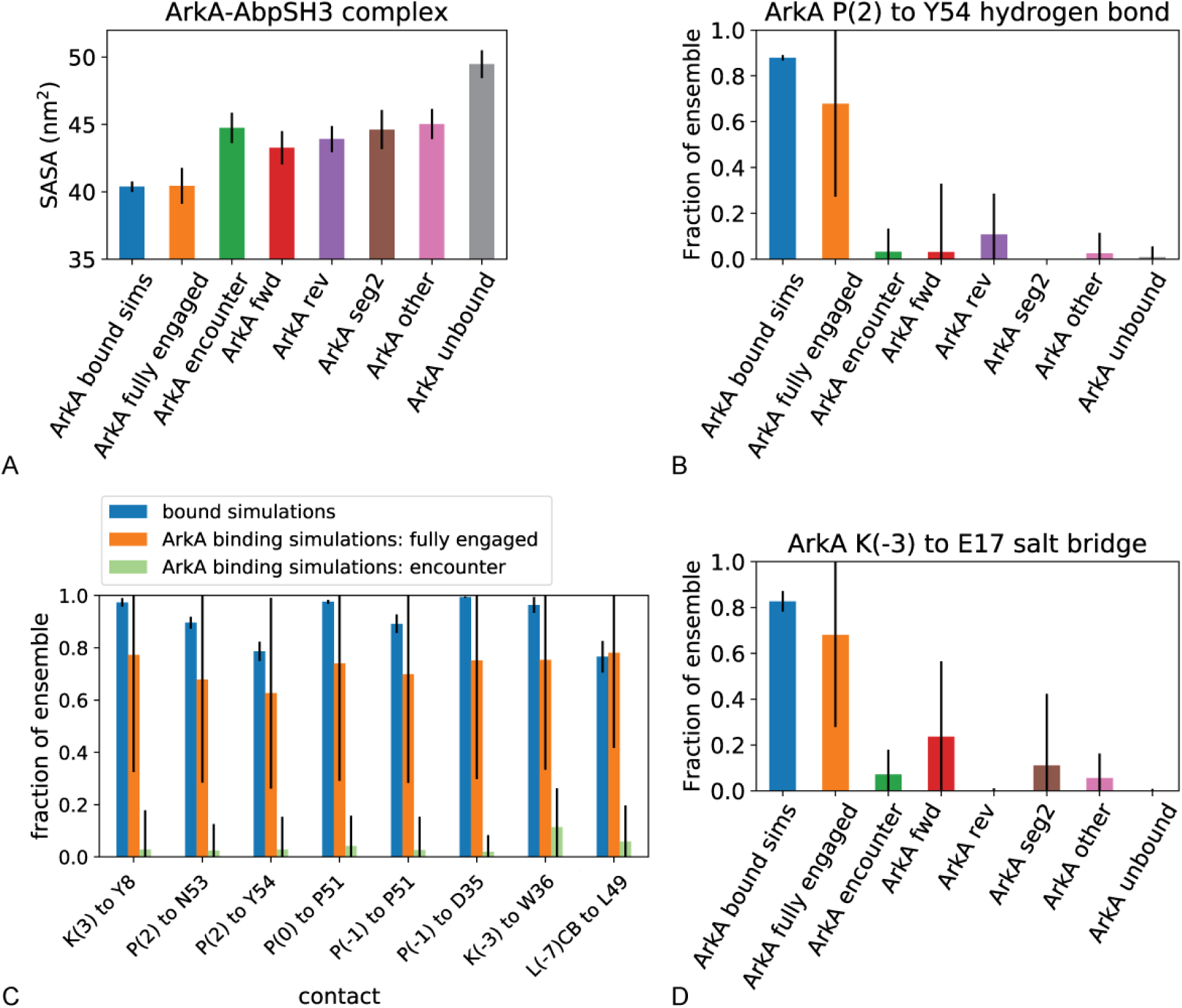
Solvent accessibility and specific intermolecular interactions for ArkA-AbpSH3 complex. A) Average solvent accessible surface area (SASA) of the ArkA-AbpSH3 system in the bound simulations (first bar) and binding simulations, by state of the complex. B) Occupancy of the P(2) to Y54 hydrogen bond in the bound simulations and ArkA binding simulations by state of the complex. C) Specific hydrophobic contacts between ArkA and the AbpSH3 binding surface in the fully engaged state and the encounter complex. D) Occupancy of the K(−3) to E17 salt bridge in the bound simulations and ArkA binding simulations by state of the complex. Error bars represent the standard deviation between independent simulations.

Additionally, in transitioning from the encounter complex to the fully engaged complex, one to two specific short-range hydrogen bond or salt bridge interactions appear (S14 Fig). In particular, in the bound simulations, there is one hydrogen bond, from the ArkA P(2) carbonyl oxygen to the AbpSH3 Y54 side chain hydroxyl group, that is present more than any others, in 88% of the simulated ensemble (Fig 10B). We also found that a short-range electrostatic salt bridge forms between ArkA K(−3) and AbpSH3 E17 in 82% of the bound simulations (Fig 10D). These specific, short-range interactions are rarely formed in the encounter ensemble, indicating that they may also to help to stabilize the fully engaged state and prevent unbinding. Previous mutation studies have found that mutating K(−3) or P(2) causes a large reduction in binding affinity of ArkA [72], possibly in part due to disruption of the salt bridge or hydrogen bond that these residues form. The K(−3) mutation had the largest effect on binding affinity [72], which may also be in part due to its specific hydrophobic interactions in the fully engaged complex (Fig 10C). The other mutation that caused a significant reduction in binding affinity was L(−7) [72], which is a hydrophobic residue that is also buried when the fully engaged complex forms in our simulations (Fig 10C), indicating the importance of these specific hydrophobic interactions. While the encounter complex is characterized by nonspecific electrostatic and hydrophobic interactions, the fully engaged complex requires more specific and complete hydrophobic contacts between ArkA and the AbpSH3 binding surface and is additionally geometrically constrained by the formation of a specific hydrogen bond and salt bridge.

### ArkA-AbpSH3 two-step binding model

Putting together all of our data from the binding simulations, we can form a picture of how ArkA binding to AbpSH3 proceeds (Fig 2). Initially the ArkA peptide is (orientationally) steered by long-range electrostatic attraction to the AbpSH3 binding surface and forms a metastable encounter complex (step 1). This encounter complex is stabilized by transient and nonnative interactions, including long-range electrostatic interactions and partially engaged hydrophobic contacts, but the binding surface is still partially solvated, especially SII. Even before formation of the encounter complex, seg1 of ArkA is pre-folded into a PPII helix, and in the encounter complex it often forms nonspecific hydrophobic interactions with SI and nonnative hydrogen bonds, although part of the peptide is still solvated in any given conformation, and the P(2) to Y54 hydrogen bond and K(−3) to E17 salt bridge are essentially absent. The seg1 PPII helix can interact with SI of AbpSH3 in either the forward or reverse orientation in the encounter complex, but the reverse orientation requires that seg2 interact with solvent rather than SII of AbpSH3. From the forward state of the encounter complex, ArkA-AbpSH3 can transition to the fully engaged state through a zippering process [5], burying hydrophobic sidechains and displacing more solvent, particularly on seg2 and SII, and forming the P(2) to Y54 hydrogen bond and K(−3) to E17 salt bridge (step 2). This transition also coincides with seg2 of ArkA becoming a bit more structured, although it is clear that the fully engaged state is still in dynamic exchange, consistent with previous co-liner chemical shift perturbation measurements [40]. The AbpSH3 binding pathway that we have characterized (Fig 2) is similar to that proposed for the c-Crk N-SH3 domain [21, 23], although our simulations provide more sampling of individual binding trajectories, allowing us to quantitatively characterize the presence of different conformational states and long and short range interactions that are present in the encounter complex ensemble. In combination with previous studies, our results indicate that this pathway may be a common binding progression for proline-rich peptides binding to SH3 domains.

### ArkA intrinsic structure affects the binding pathway

Even in the fuzzy encounter complex ensemble, seg1 of ArkA is largely folded into a PPII helix, exhibiting a binding strategy also employed by other IDPs, where one segment with pre-formed structure can dock into place first, followed by the coupled folding and binding of more flexible segments [5,7,11,25,26,48]. Polyproline sequences are especially well adapted to this strategy as PPII helices are rigid, allowing them to project from folded parts of a larger full length protein, and hydrophobic yet also highly soluble in water [5]. This involvement of pre-formed structure in the binding pathway is useful for modulating the entropy change on binding by tuning the degree of structure present in the unbound state [5, 24]. Sequence changes that change the PPII propensity but maintain the same fully-engaged SH3 complex could be a mechanism to tune the association rate, affinity, and specificity of the interaction for different cellular functions. Seg2 of ArkA is more flexible and therefore less likely to form the first tight interactions with the AbpSH3 binding surface. Modulating the amount of intrinsic 3^10^ helix structure in seg2 would be unlikely to affect the peptide binding affinity [25], since this segment also remains quite flexible in the fully engaged complex.

### Binding rates probed by NMR and MD simulations

Using NMR CPMG experiments, we determined that the ArkA peptide binds quickly, on the μs timescale, at our experimental concentrations (Table 5) [40]. In the binding simulations, ArkA generally reached a stable state in the 1 μs of simulation time, but often this state was part of the metastable encounter complex rather than the fully engaged state. Based on the 9 simulations (out of 50) that did reach the fully engaged state, we calculated a binding rate constant, *k*_on_, which we compare to the experimental binding rate (Table 5). Our simulations show that binding happens on a similar timescale to the binding rates measured by NMR. However, the *k*_on_ value from our simulations is imprecise because most simulations remained in the encounter complex (panel A in S16 Fig), and we are only able to definitively state that our simulations are not inconsistent with the rate constants determined by NMR. We can more precisely calculate a rate constant, *k*_1_, for step 1 of binding (Fig 2), since all simulations reached the encounter complex (panel B in S16 Fig). Step 1 occurs more than an order of magnitude more rapidly than the complete binding process. This extremely rapid *k*_1_ indicates that *k*_2_ could be quite low and still result in the fast *k*_on_ observed experimentally. For example, using this value of *k*_1_, a rough approximation of *k*_-1_ from our simulations (2.6 × 10^7^ s^-1^), and the experimental value of *k*_on_, we can calculate *k*_2_ based on the steady-state approximation for a two-step reaction. If we approximate *k*_-2_ = 0, *k*_on_ is given by

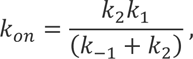

and we can solve for *k*_2_ in terms of *k*_on_:

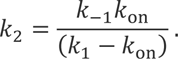

Based on this calculation, we find that *k*_2_ is 6.8 × 10^5^ s^-1^. This corresponds to a timescale for step 2 of about 1 μs, which is similar to the timescale of binding for a single ArkA molecule at our experimental SH3 domain concentration (∼8 μs).

**Table 5.** Binding rate constants for ArkA and seg1 determined from binding simulations and NMR experiments.

| | | $K_{on}$ | | $k_1$ |
| --- | --- | --- | --- | --- |
| Construct | Simulations used | Simulations ($s^{-1} M^{-1}$) | NMR ($s^{-1} M^{-1}$) | Simulations ($s^{-1} M^{-1}$) |
| ArkA | $50 \times 1 \mu s$ | $6 \times 10^7 *$ | $1.21 \pm 0.07 \times 10^8$ | $4.8 \pm 0.6 \times 10^9$ |
| seg1 | $50 \times 500 ns$ | $5 \times 10^7 *$ | Not performed | $9 \pm 1 \times 10^9$ |

\* The precision of the values of $k_{on}$ based on simulations cannot be estimated from the small number of times binding in the fully engaged state was observed.

Although our simulations clearly show a two-step binding process for ArkA-AbpSH3, previous kinetics experiments on SH3 domains have shown rapid binding consistent with a diffusion-limited process that occurs in a single step, enhanced by electrostatic steering [43]. The Fyn SH3 domain binds its peptide partner with a *k*_on_ of 1.03 × 10^8^ s^-1^ M^-1^ [38], c-Crk N-SH3 binds Sos with a *k*_on_ of 2 × 10^9^ s^-1^ M^-1^ [23], and the CrkII N-SH3 binds JNK1 with a *k*_on_ of 1.06 × 10^8^ s^-1^ M^-1^ and NS1 with a *k*_on_ of 1.1 × 10^10^ s^-1^ M^-1^ [89] (all with salt concentrations similar to or slightly lower than our experiments). However, SH3 binding peptides are IDPs, which makes the theoretical diffusion limited rate more difficult to calculate than for ordered proteins, and some IDPs seem to exceed the upper limit for binding [43, 90]. One study of a disordered region of PUMA binding to Mcl-1 found that an association rate that at first seemed to be diffusion limited in fact showed a temperature dependence for *k*_on_, indicating an energy barrier in the association process, and therefore two-step binding [43]. In our simulations, the diffusion limited association rate with an electrostatic enhancement is captured by *k*_1_; however, the overall association rate, *k*_on_, also depends on step 2 in our binding model (Fig 2).

In our simulations, step 1 (formation of the encounter complex) happens about two orders of magnitude more rapidly than the overall binding process, indicating that the transition state for binding occurs after the formation of the encounter complex. In fact, initial formation of the encounter complex appears to be a downhill process with respect to free energy, as measured along the binding distance reaction coordinate (S17 Fig). Although the step 1 rate depends on the concentration of peptide and SH3 domain, the number of transitions out of the encounter complex to either the unbound (reverse of step 1) or fully engaged state (step 2) is independent of concentration, and therefore easier to compare directly, so we focused on comparing these transitions. In our 50 independent simulations of ArkA binding, we observe 51 transitions from the encounter complex to the unbound state and only 9 from the encounter complex to the fully engaged state. This indicates that the encounter complex is approximately 5 times more likely to transition to the unbound state than to the fully engaged state. Although in our simulations the unbound state does not appear to be a stable minimum on the free energy landscape, in the cellular environment, transition to the unbound state would be part of a transition between different protein-protein interactions, such as ArkA interacting with another part of the larger Ark1p protein. Our simulations indicate that when ArkA has formed an encounter complex with AbpSH3, it is still more likely for it to unbind and begin interacting with something else than to proceed to the fully engaged state.

Our simulation result showing that the rate limiting step for ArkA binding to AbpSH3 occurs after the encounter complex formation contrasts with previous experimental data that indicate a two-state binding process [40]. Typically, when a single association rate is observed for a two-step binding reaction, it indicates that step 2 is very fast compared to step 1 [7], but this does not appear to be the case for SH3 domain binding based on this and other simulation studies where the encounter complex forms more quickly than the fully engaged state [23, 69]. In fact, in our simulations, step 1 of binding actually proceeds downhill, consistent with other studies of electrostatically enhanced binding [29, 30]. Other MD simulation studies of IDP binding have also revealed encounter complexes that form quickly, followed by a slower transition to the fully engaged complex [50,54,58]. One alternative explanation for the apparent two-state binding is that the encounter complex is only present at a very low population (< 0.5%), and therefore not detectable by NMR [38].

### The role of the encounter complex and hydrophobic interactions in binding kinetics and function

Our two-step binding model (Fig 2) that includes a fuzzy encounter complex stabilized by nonspecific hydrophobic and electrostatic interactions followed by formation of native contacts in the fully-engaged complex is observed in simulations of other IDP binding proteins, including other SH3 domains [21, 23], the PDZ domain [54], self-binding proteins [57], and the TAZ1 domain [50]. IDP complexes that lack strong charge complementarity, such as pKID and KIX, are similar, but rely mainly on nonspecific hydrophobic interactions to stabilize the encounter complex [48]. The presence of a metastable encounter complex intermediate in the binding pathway allows nonnative interactions to play an important role in the binding process. The nonspecific encounter complex can form quickly, and nonnative, transient interactions (both electrostatic and hydrophobic) allow the encounter complex to remain flexible and avoid being trapped in a state that is off-pathway for complete binding. In this binding model, hydrophobic interactions are critical in both the encounter and fully engaged complexes. While long-range electrostatic interactions only form during step 1 of binding and then remain essentially constant, hydrophobic interactions are critical to steps 1 and 2, forming nonspecifically in the transition to the encounter complex and specifically, to bury more surface area, when transitioning to the fully engaged complex. If binding only occurred in one step, mutations that affect hydrophobic interactions would only affect the dissociation rate, and not association rate of the peptide. However, with our two step binding model, we predict that hydrophobic interactions affect the stability of both the encounter complex and fully engaged state, and therefore play a role in determining the overall association rate.

The central role of hydrophobic interactions in SH3 binding was also observed by Meneses and Mittermaier [31]. They find that electrostatic rate enhancement of binding to the Fyn SH3 domain is minimal since long-range electrostatic interactions do not significantly increase the association rate compared to hydrophobic interactions. In our model (Fig 2), hydrophobic interactions form during both reaction steps and could have large effects on the association rate as well as the dissociation rate. This is consistent with the differences in CrkII N-terminal SH3 binding by the virus protein NS1 and the endogenous binding partner JNK1 observed by Shen *et al*. [89]. While the increased binding affinity and higher association rate of NS1 has been attributed to its higher positive charge [89], NS1 also contains more hydrophobic residues than JNK1, particularly within the PxxPx+ motif, which likely also has an effect on association since hydrophobic interactions enhance the formation of the encounter complex and fully engaged complex.

The encounter complex would likely play an important functional role in SH3 binding in the cellular context. Competition between binding partners may need to be tuned by modulating the encounter complex to determine which interaction will be dominant, as in the case of CITED2 competing with HIF-1α to bind TAZ1 [91]. CITED2 forms an encounter complex with TAZ1 while HIF-1α is bound, which allows it to completely displace HIF-1α even though both partners have similar affinities to TAZ1. There is evidence that AbpSH3 can form an intramolecular interaction with a proline-rich sequence of the Abp1p protein, which may inhibit binding of other partners [92]. The formation of an encounter complex in this case, may allow the competition between binding partners to be tuned as necessary for function, instead of being dictated only by their relative affinities. The transient interactions of the SH3 encounter complex may also be critical when SH3 domains need to locate a small proline-rich binding motif within a longer disordered protein segment. The domain could rapidly associate with (positively charged) disordered sequences, while the interaction remains fuzzy enough to allow rearrangement or even translation of the SH3 domain along large stretches of disordered sequence until the specific PPII partner segment can align with the binding surface and lock into the fully engaged state. This search process may resemble a transcription factor searching an elongated DNA strand for its promotor site. Thus, the encounter complex we have characterized in this study is likely an important functional intermediate in the binding pathway of this ubiquitous protein interaction domain.

## Conclusions

Through examination of ArkA-AbpSH3 binding by MD simulations and NMR experiments, we have created a two-step binding model that includes the formation of a heterogeneous encounter complex stabilized by transient, nonspecific hydrophobic and electrostatic interactions. ArkA samples many states with different degrees of folding and binding before reaching the fully engaged state, though contacts with the domain are limited to the canonical highly acidic binding surface. The fuzziness of the encounter complex ensemble allows multiple paths to the fully engaged state, driven by specific hydrophobic interactions and a key hydrogen bond and salt bridge rather than long-range electrostatics. The PxxP motif in seg1 is preformed in a PPII helix, which locks into the hydrophobic grooves of SI in a zippering mechanism during step 2 of binding. The more disordered seg2 may prevent the peptide from binding in reverse as a result of the pseudo-palindromic PxxP motif in seg1. While encounter complex formation is diffusion limited and enhanced by electrostatic orientational steering, the transition to the fully engaged state is slower and relies on specific hydrophobic interactions and short-range electrostatic interactions. In the cellular context, rapid formation of the encounter complex stabilized by transient, nonspecific interactions could allow SH3 domains to search for proline-rich motifs within disordered sequences. Our binding model and encounter complex characterization give insights into the mechanism that SH3 domains use to perform a wide variety of functions. Future studies of AbpSH3 thermodynamics and kinetics would further elucidate the different roles for electrostatic and hydrophobic interactions in the binding pathway.

## Supporting information

Combined pdf for SI text, figures, and captions

Readme for data

## Acknowledgements

The authors thank Michael Latham at Texas Tech University for help processing the CPMG data and Kristina Foley for important preliminary simulations and analysis. K.A.B. thanks Michael Donnelly for computational support. K.A.B. thanks the MERCURY Consortium for mentoring support.

## S1 Text. Seg1 binding results

**S1 Table. Temperatures (in Kelvin) used in the unbound ArkA replica exchange simulations.**

**S2 Table. Summary of volume, concentration, and binding frequency for the two binding simulations.**

**S3 Table. Summary of water box dimensions for each simulated system.**

**S1 Fig. Pairwise distances used in the determination of whether the encounter simulation has reached the bound state.** The pairs in red and magenta where used for both the seg1 and ArkA1 simulations and the cyan pairs where added for the ArkA simulations. The distances were determined based on the SH3 residues whose chemical shifts where used to determine binding in NMR experiments. The pairwise binding surface distance in the NMR structures ranges from 7.25 to 7.62 Å.

**S2 Fig. Running averages of measures used to determine convergence of REMD simulations.** ArkA structural measures plotted vs. simulation time for each of the independent REMD simulations. The first 50 ns of each independent simulation was removed before analysis.

**S3 Fig. Representative autocorrelation of replica state index graph, for one simulation, showing that the replica exchange was exchanging as expected and the number of replicas was sufficient.**

**S4 Fig. Overlay of the 20 ArkA conformations from the NMR (2RPN) ensemble with the SH3 domain aligned** [72].

**S5 Fig. Percentage of time each ArkA residue is spending in PPII Helix or 3-10 Helix in the NMR ensemble (2RPN)** [72].

**S6 Fig. Binding surface distance and dihedral angle RMSD for NMR structures (2RPN)** [72]. The distance between ArkA and the binding surface of AbpSH3 is graphed against the dihedral angle RMSD for the bound simulations with the 20 NMR structures shown as red x’s. The NMR structures all have low binding surface distances, but cover a range of different dihedral angle RMSD values. Darker shading indicates a larger fraction of the total bound simulation ensemble, as indicated by the color bar.

**S7 Fig. Contact maps of the fully engaged state from the NMR ensemble (2RPN) and bound simulations.** The darker squares indicate more of the ensemble with that contact. The red and blue squares along the AbpSH3 residue index show which amino acids are in SI and SII, respectively. The black line indicates the separation of seg1 and seg2, and the single letter amino acid codes are included for ArkA and the residues in SI and SII.

**S8 Fig. Starting structures for ArkA binding simulations.** The conformational ensemble of unbound ArkA from the REMD simulations is plotted in terms of end-to-end distance and dihedral angle RMSD, with starting structures for ArkA binding simulations indicated by blue circles. End-to-end distance is the distance between the C and N-terminal ends of ArkA and dihedral angle RMSD is calculated only for ArkA with the lowest energy NMR structure (2RPN) as the reference [72].

**S9 Fig. Histogram of dihedral RMSD based on only the seg1 dihedral angles from the unbound simulation ensemble.** The vertical line at 38.1° indicates the cutoff that was determined between the two states (native and nonnative conformations).

**S10 Fig. Histogram of dihedral RMSD for the full ArkA peptide from the unbound simulation ensemble.** The vertical line at 33.7° indicates the cutoff that was determined between the native folded and nonnative states.

**S11 Fig. Distance between seg1 and the binding surface of AbpSH3 over time for an example seg1 binding simulation.** The black lines correspond to our definition of the encounter complex (23 Å) and the fully engage complex (11.5 Å).

**S12 Fig. Distance between seg1 and the binding surface of AbpSH3 graphed against the dihedral angle RMSD for seg1 binding simulations.** Darker shading indicates a larger fraction of the total ensemble, as indicated by the color bar. Colored boxes partition the ensemble into four states: folded and fully engaged (blue), unfolded and fully engaged (green), folded and encounter (red), unfolded and encounter (cyan).

**S13 Fig. Solvent accessible surface area (A) and occupancy of the P(2) to Y54 hydrogen bond (B) for the seg1-AbpSH3 complex in different states.** The first bar on the plot represents the solvent accessible surface area in bound simulations. Error bars represent the standard deviation between independent simulations.

**S14 Fig. Average number of hydrogen bonds (or salt bridges) between AbpSH3 and the ArkA peptide for ArkA (A) and seg1 (B) in the bound simulations (first bar) and binding simulations by state of the complex.** Error bars represent the standard deviation between independent simulations.

**S15 Fig. Specific hydrophobic contacts between the seg1 peptide and the AbpSH3 binding surface in the fully engaged and encounter complexes.** Hydrophobic contacts were selected from those hydrocarbon groups that are closest together in the NMR structural ensemble (2RPN) [72]. Contacts were defined based on a 6 Å cutoff distance between hydrocarbon groups. Error bars represent the standard deviation between independent simulations.

**S16 Fig. Fit of TCDF curve (red line) to time to formation data (blue circles) for the fully engaged ArkA complex (A) and for the ArkA encounter complex (B) from ArkA binding simulations.** These fits were used to determine *k*_on_ and *k*_1_ respectively.

**S17 Fig. Population histogram of binding surface distance from ArkA binding simulaitons.** States left of the vertical line at 11.5 Å are classified as fully engaged. States between the vertical line at 11.5 Å and the one at 23 Å are classified as the encounter complex. States right of the vertical line at 23 Å are classified as unbound. There is a clear population decrease between the fully engaged and encounter complexes, indicating a free energy barrier. It is important to note that these binding simulations do not fully sample the fully engaged state or the barrier between fully engaged and encounter, so this histogram cannot be considered an equilibrium ensemble. There is no clear barrier between the unbound state and encounter complex, indicating that formation of the encounter complex from the unbound state is downhill in free energy. Within the encounter complex, the binding surface distance reaction coordinate reveals two populations. The population between 11.5 and 15 Å is 32% of the entire encounter complex ensemble and contains almost all of the forward encounter states (67.1% other, 28.9% forward, 4.0% segment 2 only). The population between 15 and 23 Å is 68% of the encounter complex ensemble and contains all of the reverse encounter states (82.7% other, 12.8% reverse, 2.8% segment 2 only, 0.6% forward).

**S18 Fig. Fraction of frames with intermolecular contact between ArkA and AbpSH3 by binding surface distance.** The blue bars represent the fraction of frames that have at least one intermolecular contacts between ArkA and AbpSH3 at each bin along the binding surface distance reaction coordinate. The vertical black line at 11.5 Å represents the division between the fully engaged and encounter complex states, while the vertical black line at 23 Å represent the division between the encounter complex and the unbound state. While 100% of the fully engaged structures and 99.9% of the encounter complex structures have at least one intermolecular contact between ArkA and AbpSH3, 34% of the unbound structures have no contacts between ArkA and AbpSH3.

**S19 Fig. Occupancy of the K(−3) to E17 salt bridge for the seg1-AbpSH3 complex in different states.** The first bar on the plot represents the salt bridge occupancy in bound simulations. Error bars represent the standard deviation between independent simulations.

**S20 Fig. NMR ^15^N CPMG relaxation dispersion data for the amide signals of 21 AbpSH3 residues at 500 (triangles) and 800 (squares) MHz.** The top of each plot is labeled with the residue number, and 36s and 37s refer to the tryptophan sidechain NH groups.

**S1 Movie. Example simulation of initial ArkA binding to the AbpSH3 domain.**

**S2 Movie. Example simulation of initial ArkA binding to the AbpSH3 domain.**

**S3 Movie. Simulation of ArkA changing orientation from forward to reverse in the encounter complex.**

