## Supplementary material for "A disordered encounter complex is central to the yeast Abp1p SH3 domain binding pathway": Combined pdf for SI text, figures, and captions

**S1 Text. Seg1 binding results.**

Seg1 contains the class II PxxPx+ motif, which is almost the reverse of the class I +xxPxxP motif. With this pseudo-palandromic motif, seg1 is likely to be able to bind in two different orientations, and experimental data shows that seg1 binds the domain with a lower affinity than ArkA [38]. We wanted to test whether the binding pathway differs for the shorter peptide in order to examine the role of seg1 and seg2 in binding. To initiate the seg1 binding simulations, we divided the unbound ArkA ensemble into two states based on the seg1 dihedral RMSD with a cutoff of 38.1˚ (S9 Fig) and ran 25 independent seg1 binding simulations for 500 ns each, starting from each of the two states (Table 1). We found that seg1 also occupies an encounter complex before reaching the fully engaged state, but it spends more time in both forward and reverse orientations within the encounter complex ensemble than the ArkA peptide does. It may be that the short seg1 peptide is not a good proxy for how seg1 behaves as part of the longer peptide.

In the seg1 simulations, the encounter complex was ~10 times more likely to transition to the unbound state than the fully engaged state (we observed 49 transitions to unbound and only 4 to fully engaged from the encounter complex). ArkA is only ~5 times more likely to transition to the unbound state than the fully engaged state when in the encounter complex. This indicates that the barrier between unbound and the encounter complex is lower for seg1 than for ArkA, which is supported by the higher $k_1$ rate constant for seg1 in our simulations. Since the seg1 peptide has fewer groups that can interact nonspecifically with the domain in the encounter complex ensemble, it is easier for it to transition to a state where it will dissociate completely.

**S1 Table. Temperatures (in Kelvin) used in the unbound ArkA replica exchange simulations.**

The temperature used for analysis is in bold.

| | | |
|---|---|---|
| 290.00 | 330.30 | 376.20 |
| 292.37 | 332.99 | 379.27 |
| 294.76 | 335.71 | 382.36 |
| 297.16 | 338.45 | 385.49 |
| **299.59** | 341.22 | 388.63 |
| 302.03 | 344.00 | 391.81 |
| 304.50 | 346.81 | 395.01 |
| 306.99 | 349.65 | 398.23 |
| 309.49 | 352.50 | 401.48 |
| 312.02 | 355.38 | 404.76 |
| 314.57 | 358.28 | 408.07 |
| 317.14 | 361.21 | 411.40 |
| 319.73 | 364.15 | 414.76 |
| 322.34 | 367.13 | 418.14 |
| 324.97 | 370.13 | 421.56 |
| 327.62 | 373.15 | 425.00 |

**S2 Table. Summary of volume, concentration, and binding frequency for the two binding simulations.**

| | Volume (Å$^3$) | Concentration (mM) | Binding frequency ($s^{-1}$) |
|---|---|---|---|
| ArkA | 437,360 | 3.80 | $2.4 \times 10^5$ |
| seg1 | 403,200 | 4.12 | $1.9 \times 10^5$ |

**S3 Table. Summary of water box dimensions for each simulated system.**

| | Dimensions (Å) | | |
|---|---|---|---|
| Bound simulations | 49 | 49 | 49 |
| Unbound (extended) | 63 | 63 | 63 |
| Unbound (NMR) | 59 | 59 | 59 |
| ArkA binding | 71 | 77 | 80 |
| Seg1 binding | 60 | 80 | 84 |

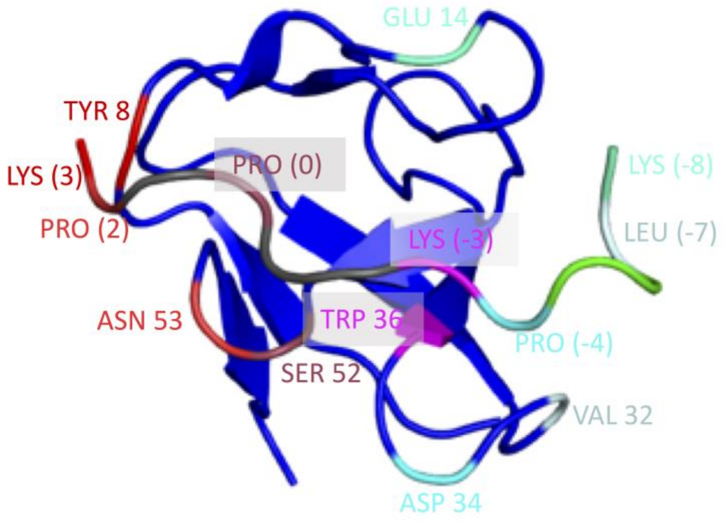


**S1 Fig. Pairwise distances used in the determination of whether the encounter simulation has reached the bound state.** The pairs in red and magenta where used for both the seg1 and ArkA1 simulations and the cyan pairs where added for the ArkA simulations. The distances were determined based on the SH3 residues whose chemical shifts where used to determine binding in NMR experiments. The pairwise binding surface distance in the NMR structures ranges from 7.25 to 7.62 Å.

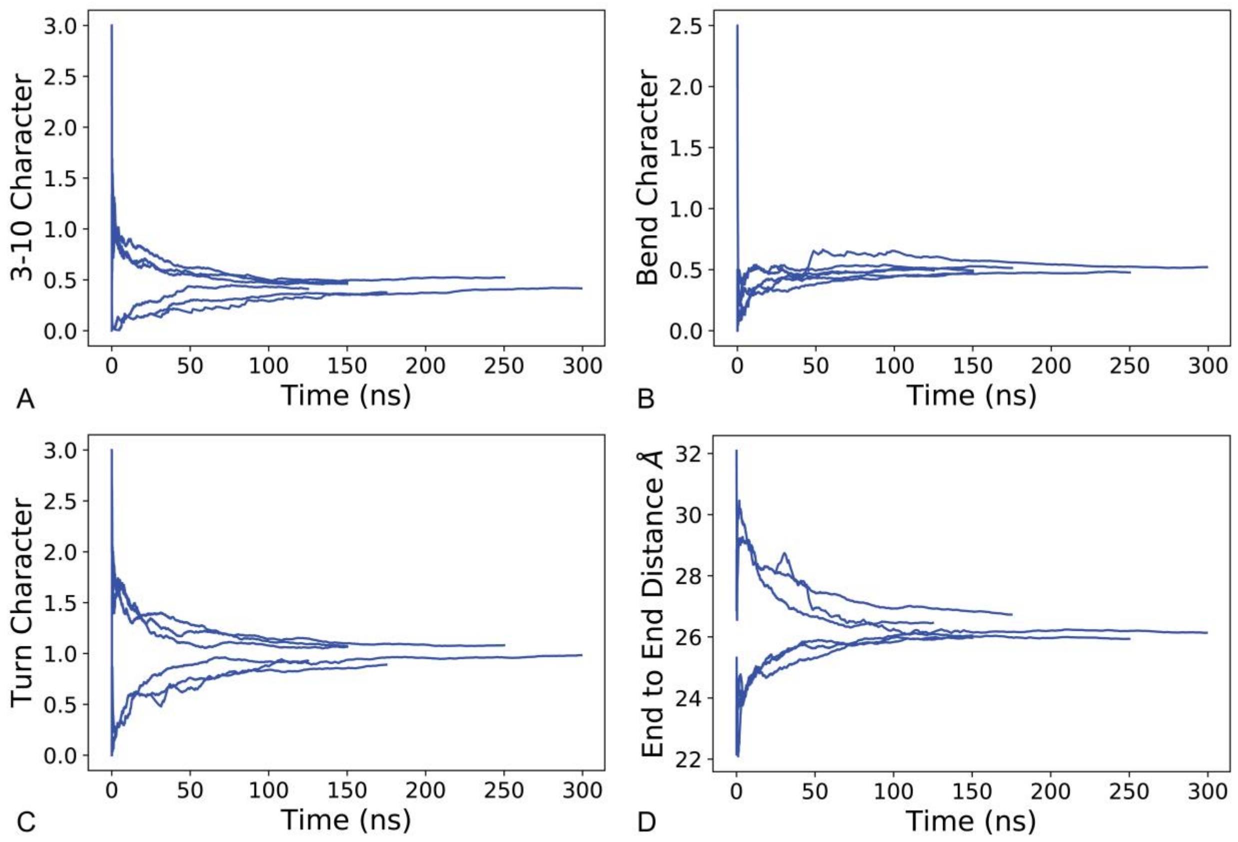


**S2 Fig. Running averages of measures used to determine convergence of REMD simulations.** ArkA structural measures plotted vs. simulation time for each of the independent REMD simulations. The first 50 ns of each independent simulation was removed before analysis.

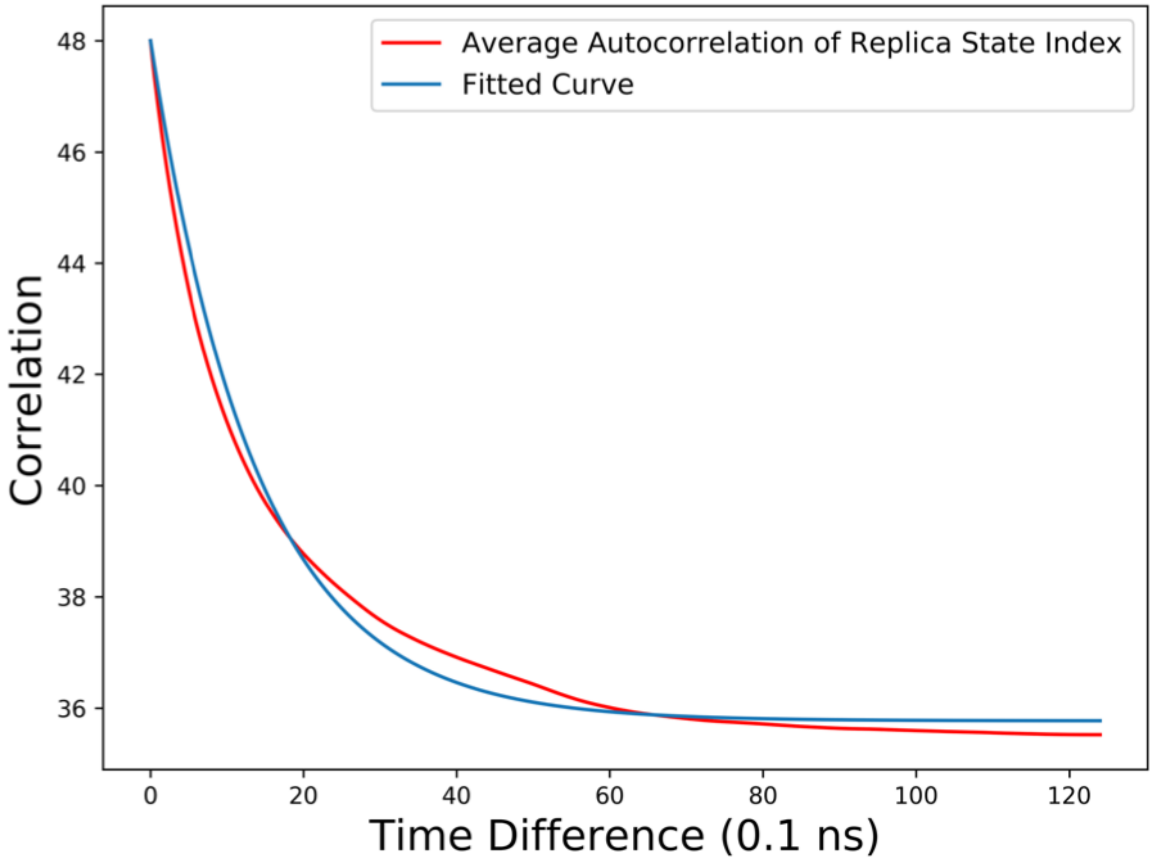


**S3 Fig. Representative autocorrelation of replica state index graph, for one simulation, showing that the replica exchange was exchanging as expected and the number of replicas was sufficient.**

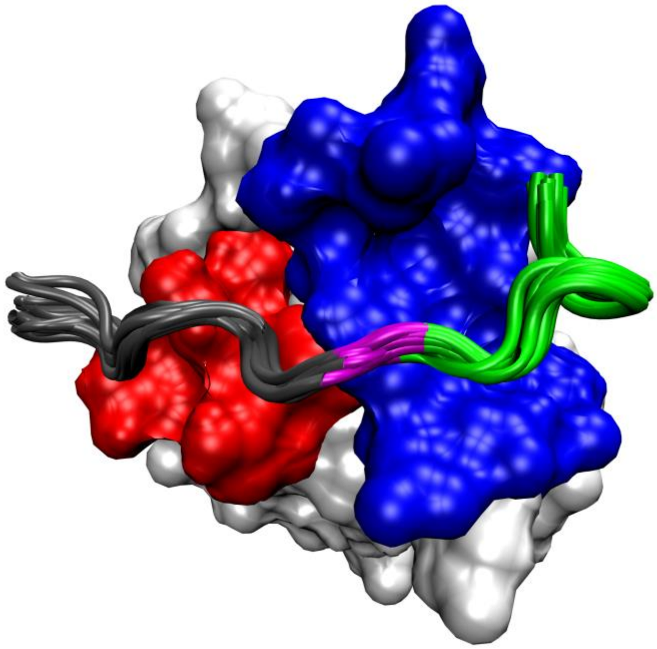

**S4 Fig. Overlay of the 20 ArkA conformations from the NMR (2RPN) ensemble with the SH3 domain aligned [69].**

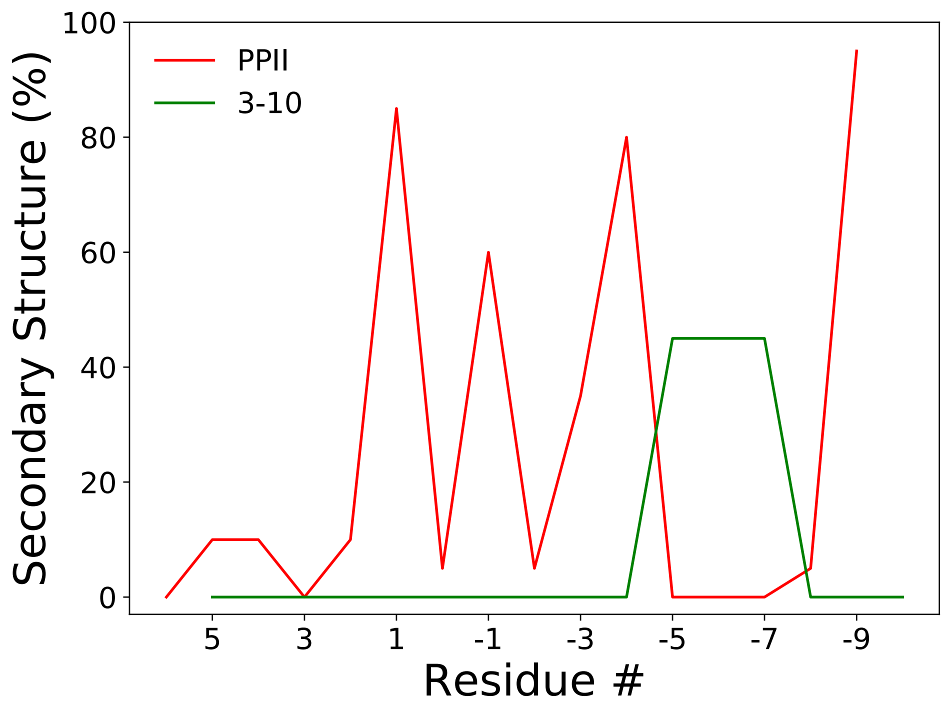


**S5 Fig. Percentage of time each ArkA residue is spending in PPII Helix or 3-10 Helix in the NMR ensemble (2RPN) [69].**

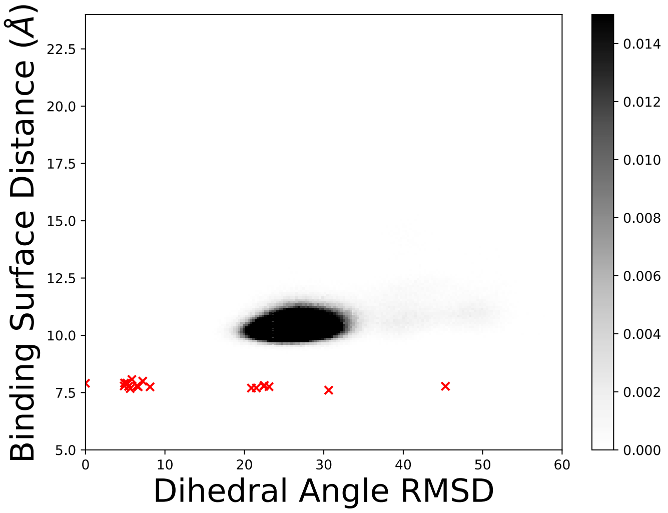


**S6 Fig. Binding surface distance and dihedral angle RMSD for NMR structures (2RPN) [69].** The distance between ArkA and the binding surface of AbpSH3 is graphed against the dihedral angle RMSD for the bound simulations with the 20 NMR structures shown as red x's. The NMR structures all have low binding surface distances, but cover a range of different dihedral angle RMSD values. Darker shading indicates a larger fraction of the total bound simulation ensemble, as indicated by the color bar.

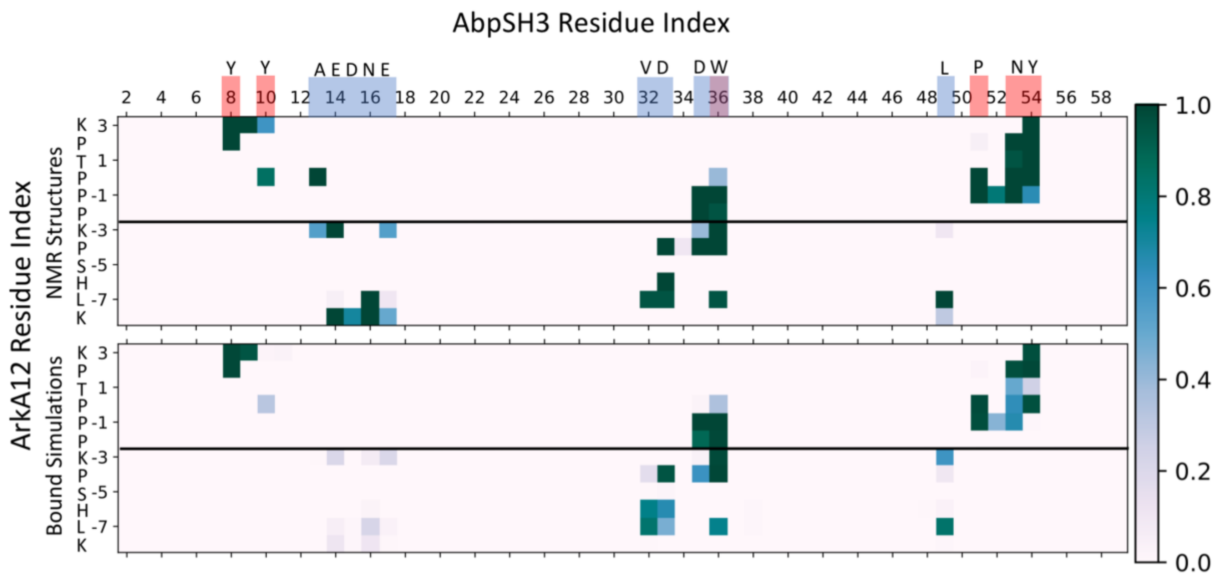


**S7 Fig. Contact maps of the fully engaged state from the NMR ensemble (2RPN) and bound simulations.** The darker squares indicate more of the ensemble with that contact. The red and blue squares along the AbpSH3 residue index show which amino acids are in SI and SII, respectfully. The black line indicates the separation of seg1 and seg2, and the single letter amino acid codes are included for ArkA and the residues in SI and SII.

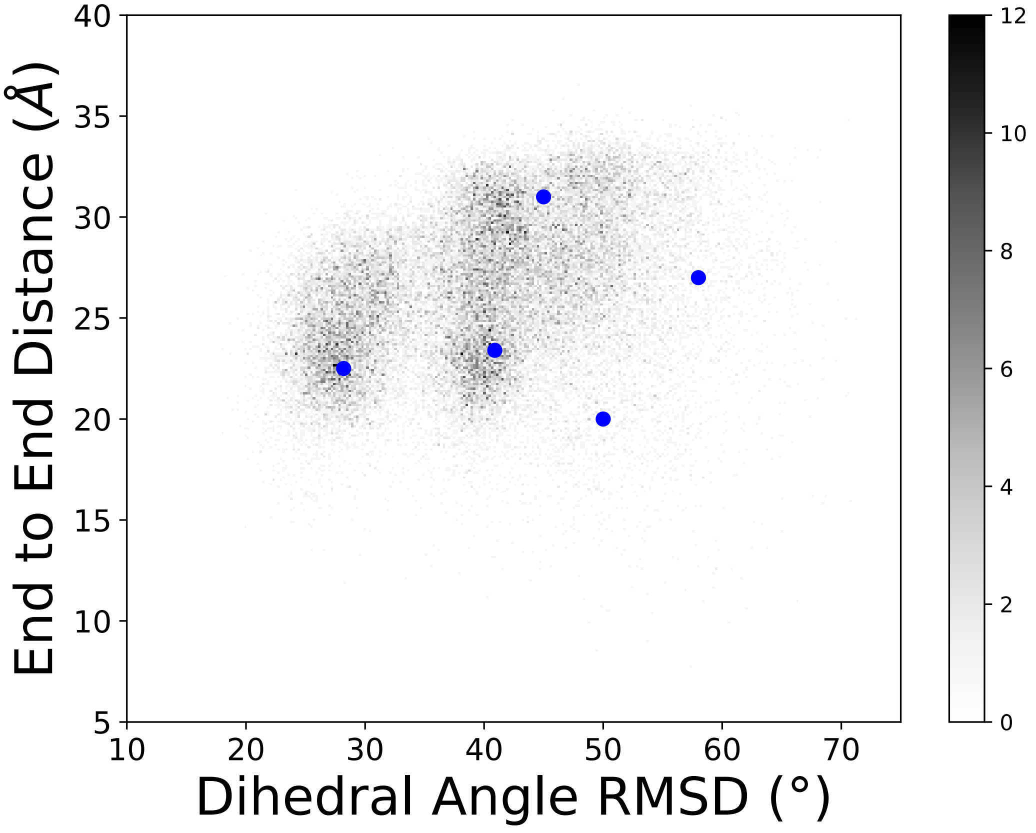


**S8 Fig. Starting structures for ArkA binding simulations.** The conformational ensemble of unbound ArkA from the REMD simulations is plotted in terms of end-to-end distance and dihedral angle RMSD, with starting structures for ArkA binding simulations indicated by blue circles. End-to-end distance is the distance between the C and N-terminal ends of ArkA and dihedral angle RMSD is calculated only for ArkA with the lowest energy NMR structure (2RPN) as the reference [69].

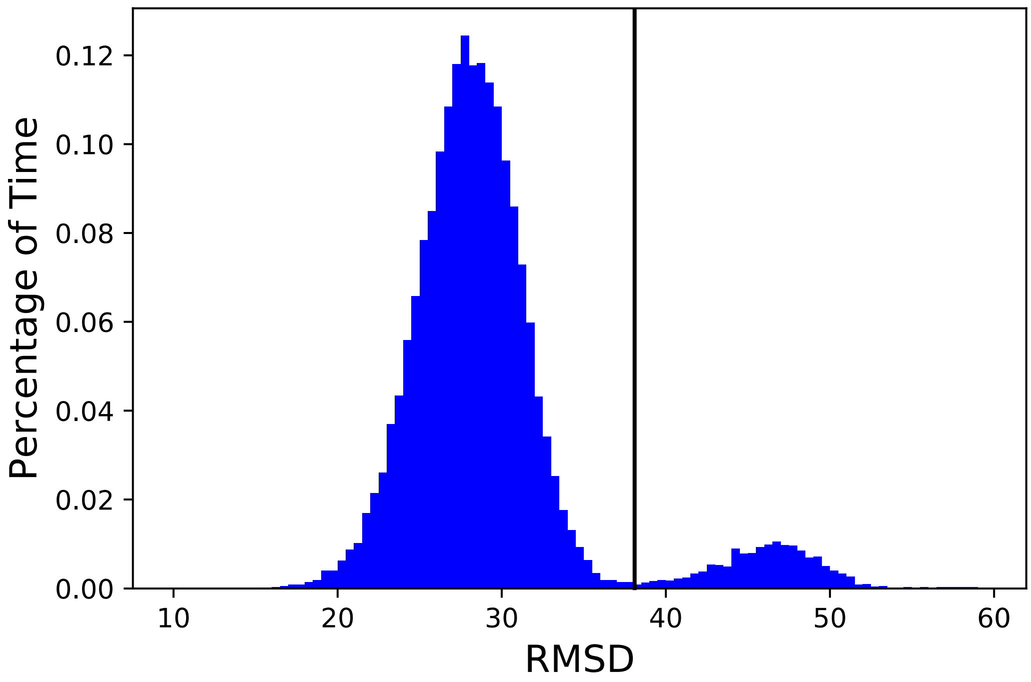


**S9 Fig. Histogram of dihedral RMSD based on only the seg1 dihedral angles from the unbound simulation ensemble.** The vertical line at 38.1° indicates the cutoff that was determined between the two states (native and nonnative conformations).

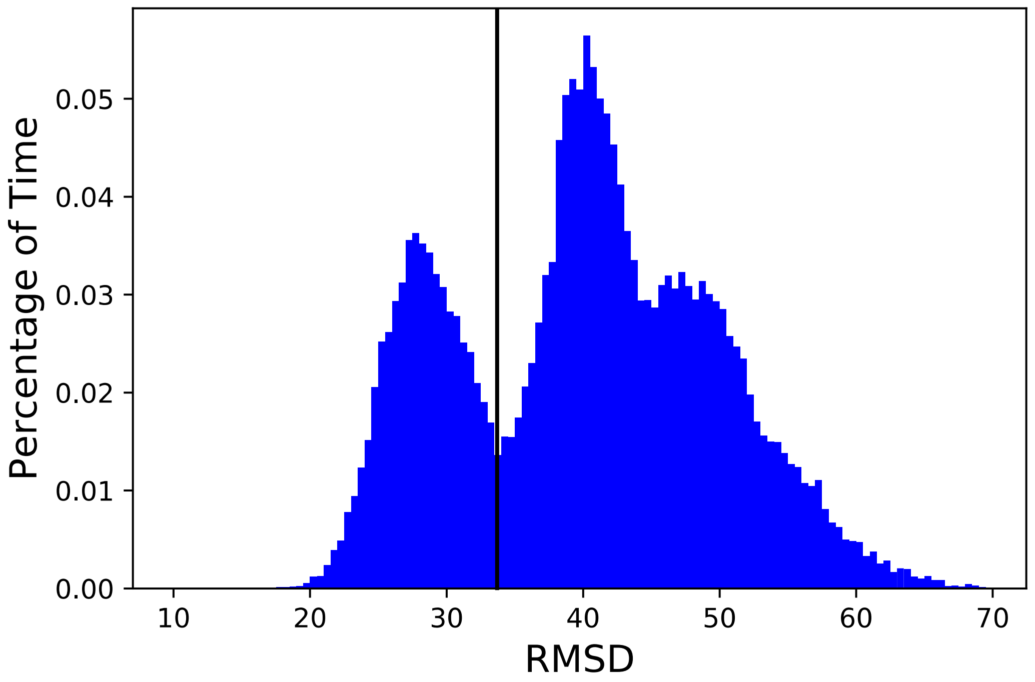


**S10 Fig. Histogram of dihedral RMSD for the full ArkA peptide from the unbound simulation ensemble.** The vertical line at 33.7° indicates the cutoff that was determined between the native folded and nonnative states.

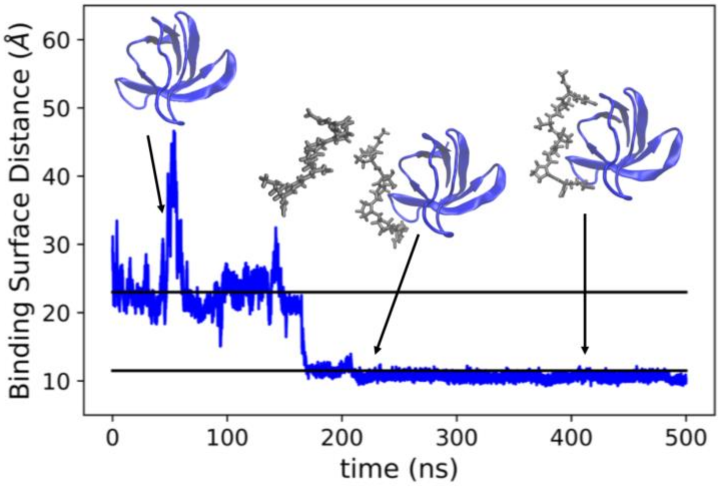


**S11 Fig. Distance between seg1 and the binding surface of AbpSH3 over time for an example seg1 binding simulation.** The black lines correspond to our definition of the encounter complex (23 Å) and the fully engage complex (11.5 Å).

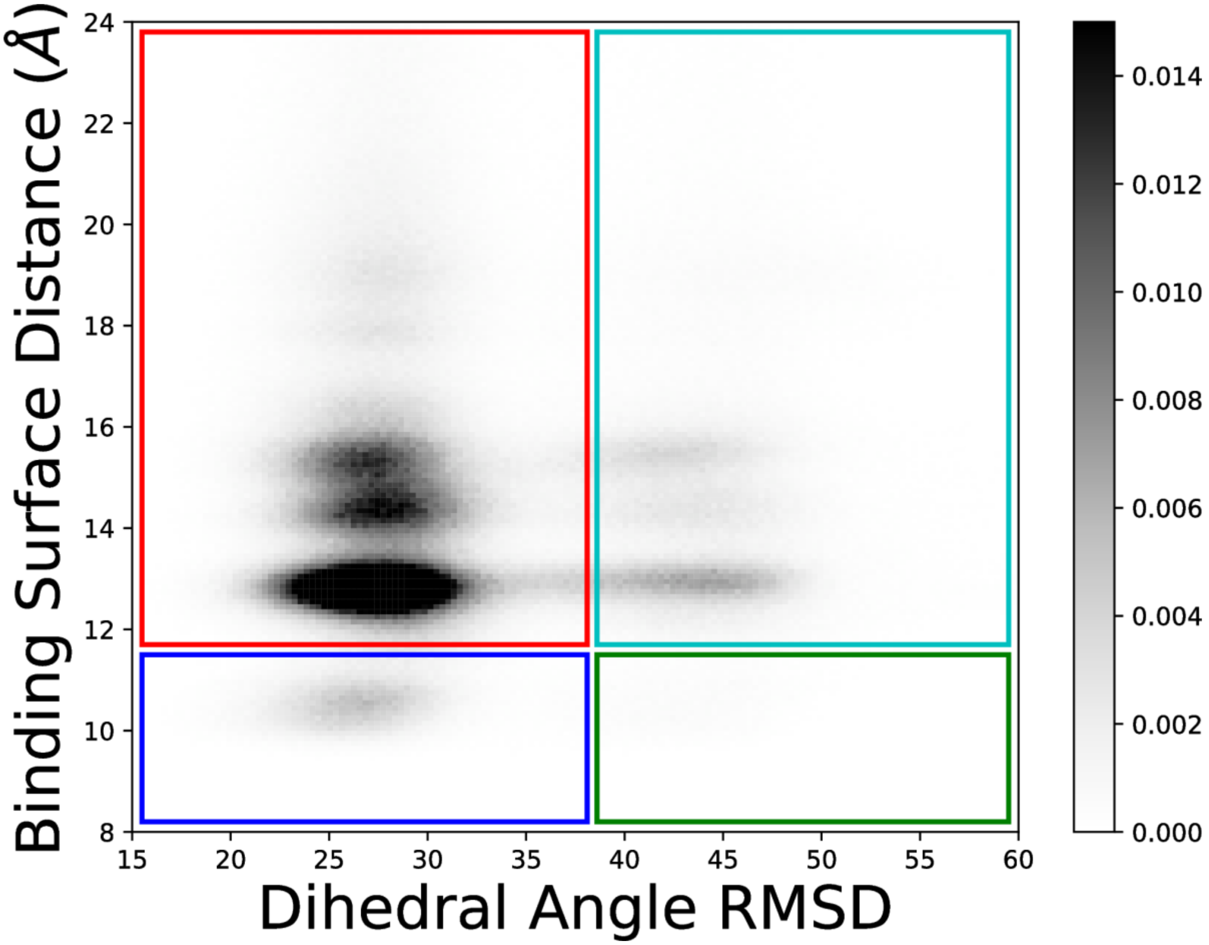


**S12 Fig. Distance between seg1 and the binding surface of AbpSH3 graphed against the dihedral angle RMSD for seg1 binding simulations.** Darker shading indicates a larger fraction of the total ensemble, as indicated by the color bar. Colored boxes partition the ensemble into four states: folded and fully engaged (blue), unfolded and fully engaged (green), folded and encounter (red), unfolded and encounter (cyan).

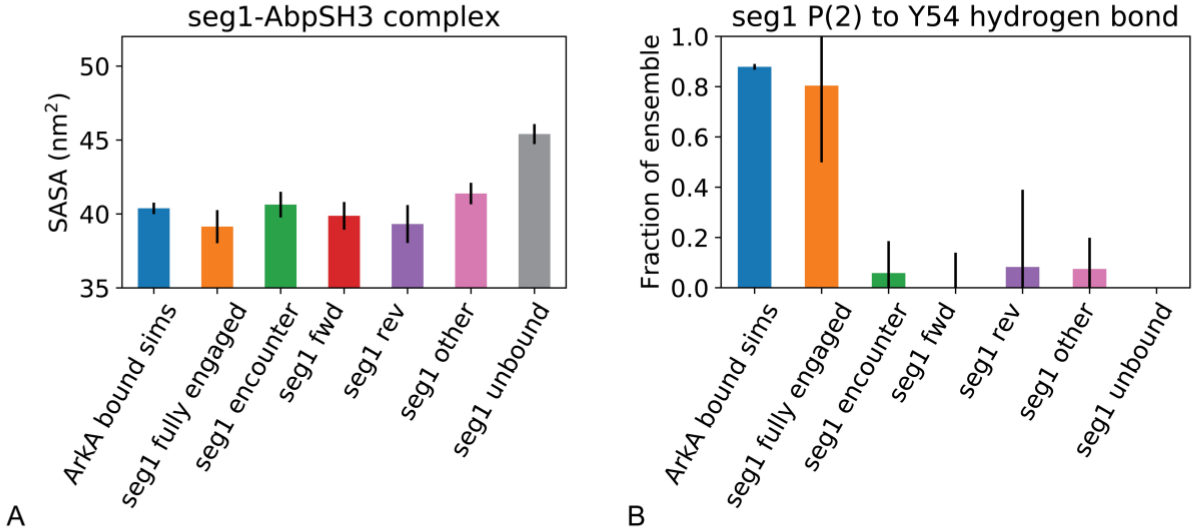


**S13 Fig. Solvent accessible surface area (A) and occupancy of the P(2) to Y54 hydrogen bond (B) for the seg1-AbpSH3 complex in different states.** The first bar on the plot represents the solvent accessible surface area in bound simulations. Error bars represent the standard deviation between independent simulations.

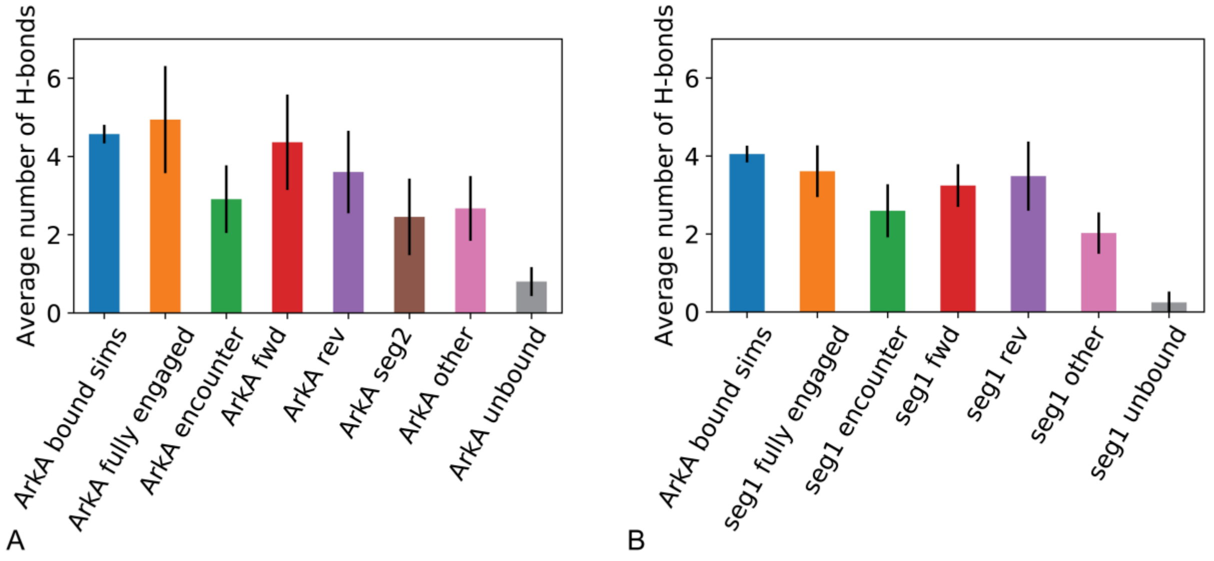


**S14 Fig. Average number of hydrogen bonds (or salt bridges) between AbpSH3 and the ArkA peptide for ArkA (A) and seg1 (B) in the bound simulations (first bar) and binding simulations by state of the complex.** Error bars represent the standard deviation between independent simulations.

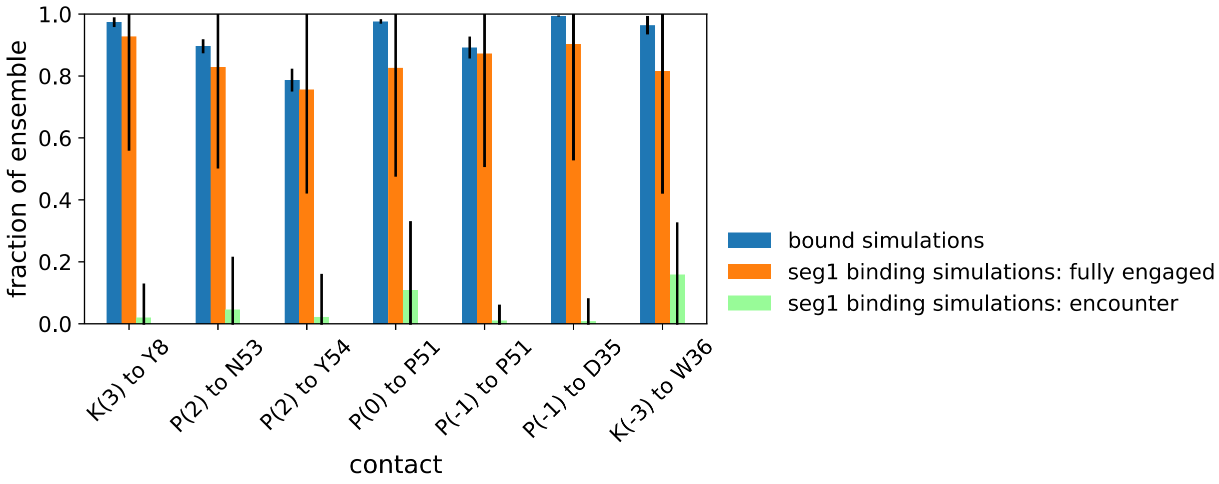


**S15 Fig. Specific hydrophobic contacts between the seg1 peptide and the AbpSH3 binding surface in the fully engaged and encounter complexes.** Hydrophobic contacts were selected from those hydrocarbon groups that are closest together in the NMR structural ensemble (2RPN) [69]. Contacts were defined based on a 6 Å cutoff distance between hydrocarbon groups. Error bars represent the standard deviation between independent simulations.

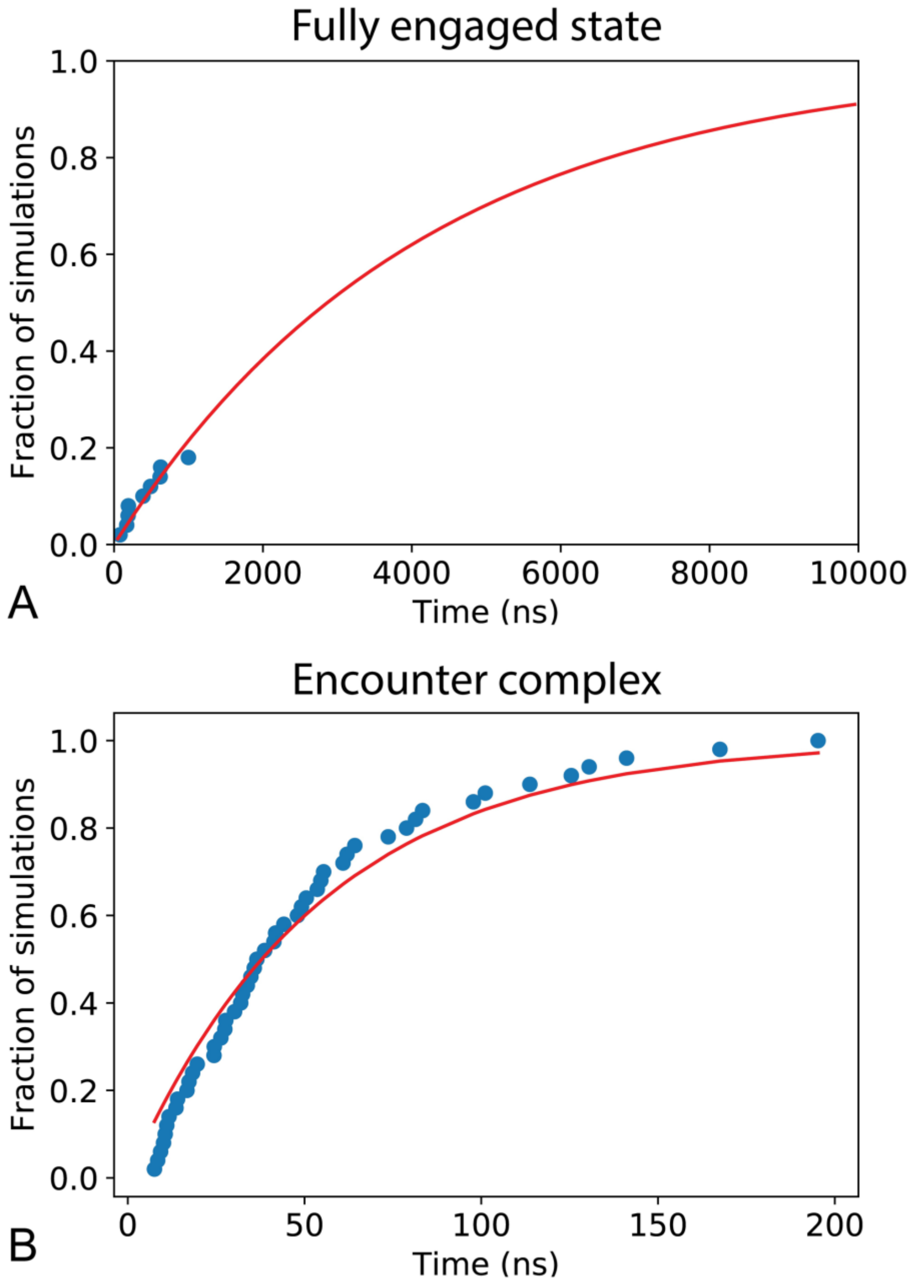


**S16 Fig. Fit of TCDF curve (red line) to time to formation data (blue circles) for the fully engaged ArkA complex (A) and for the ArkA encounter complex (B) from ArkA binding simulations.** These fits were used to determine $k_{on}$ and $k_1$ respectively.

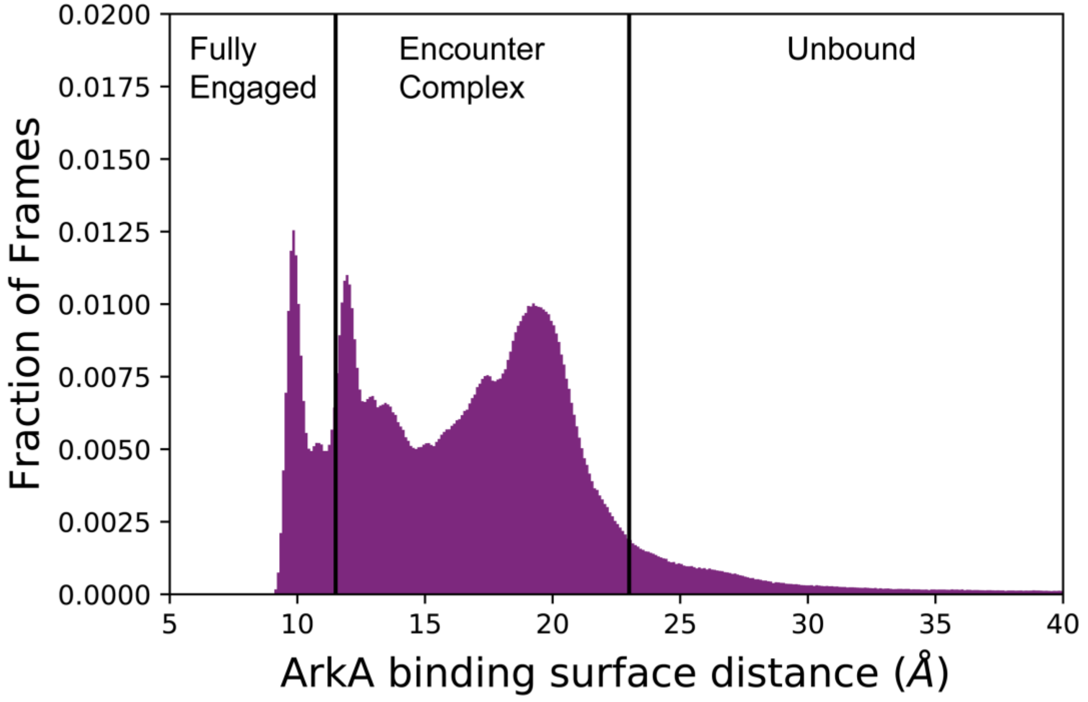


**S17 Fig. Population histogram of binding surface distance from ArkA binding simulaitons.** States left of the vertical line at 11.5 Å are classified as fully engaged. States between the vertical line at 11.5 Å and the one at 23 Å are classified as the encounter complex. States right of the vertical line at 23 Å are classified as unbound. There is a clear population decrease between the fully engaged and encounter complexes, indicating a free energy barrier. It is important to note that these binding simulations do not fully sample the fully engaged state or the barrier between fully engaged and encounter, so this histogram cannot be considered an equilibrium ensemble. There is no clear barrier between the unbound state and encounter complex, indicating that formation of the encounter complex from the unbound state is downhill in free energy. Within the encounter complex, the binding surface distance reaction coordinate reveals two populations. The population between 11.5 and 15 Å is 32% of the entire encounter complex ensemble and contains almost all of the forward encounter states (67.1% other, 28.9% forward, 4.0% segment 2 only). The population between 15 and 23 Å is 68% of the encounter complex ensemble and contains all of the reverse encounter states (82.7% other, 12.8% reverse, 2.8% segment 2 only, 0.6% forward).

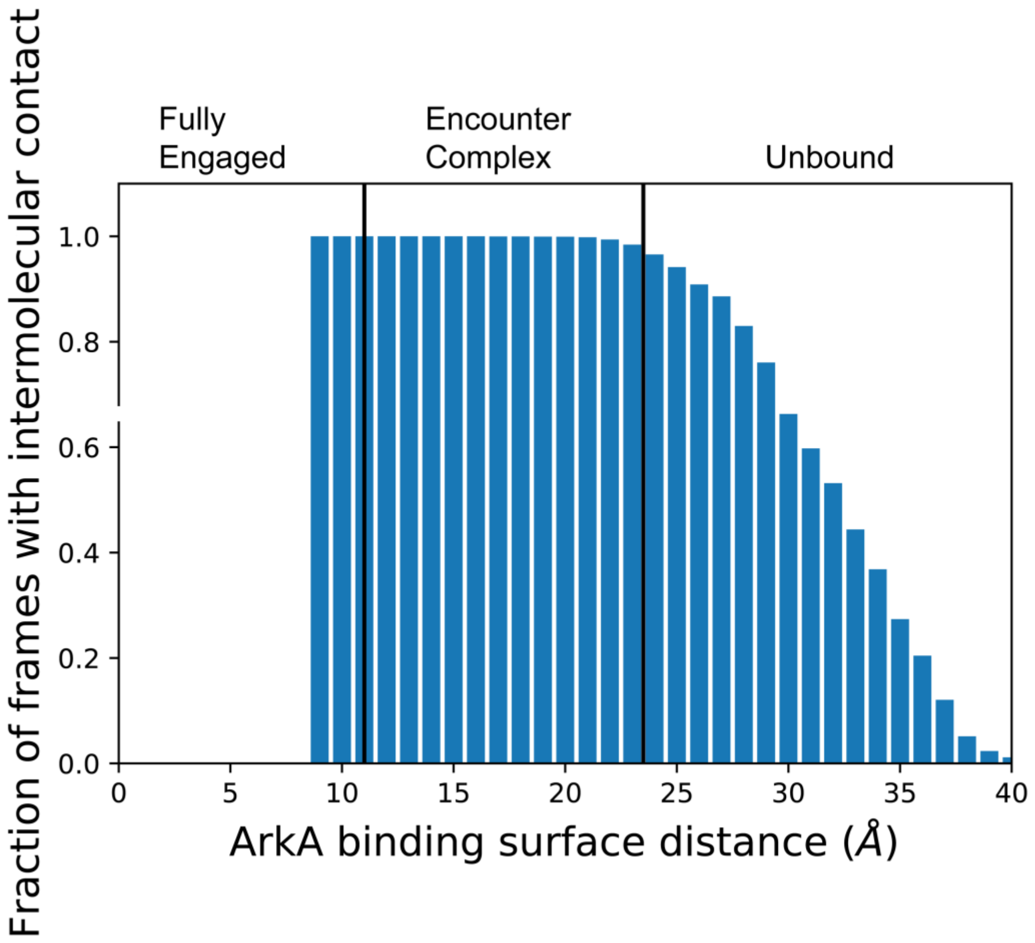


**S18 Fig. Fraction of frames with intermolecular contact between ArkA and AbpSH3 by binding surface distance.** The blue bars represent the fraction of frames that have at least one intermolecular contacts between ArkA and AbpSH3 at each bin along the binding surface distance reaction coordinate. The vertical black line at 11.5 Å represents the division between the fully engaged and encounter complex states, while the vertical black line at 23 Å represent the division between the encounter complex and the unbound state. While 100% of the fully engaged structures and 99.9% of the encounter complex structures have at least one intermolecular contact between ArkA and AbpSH3, 34% of the unbound structures have no contacts between ArkA and AbpSH3.

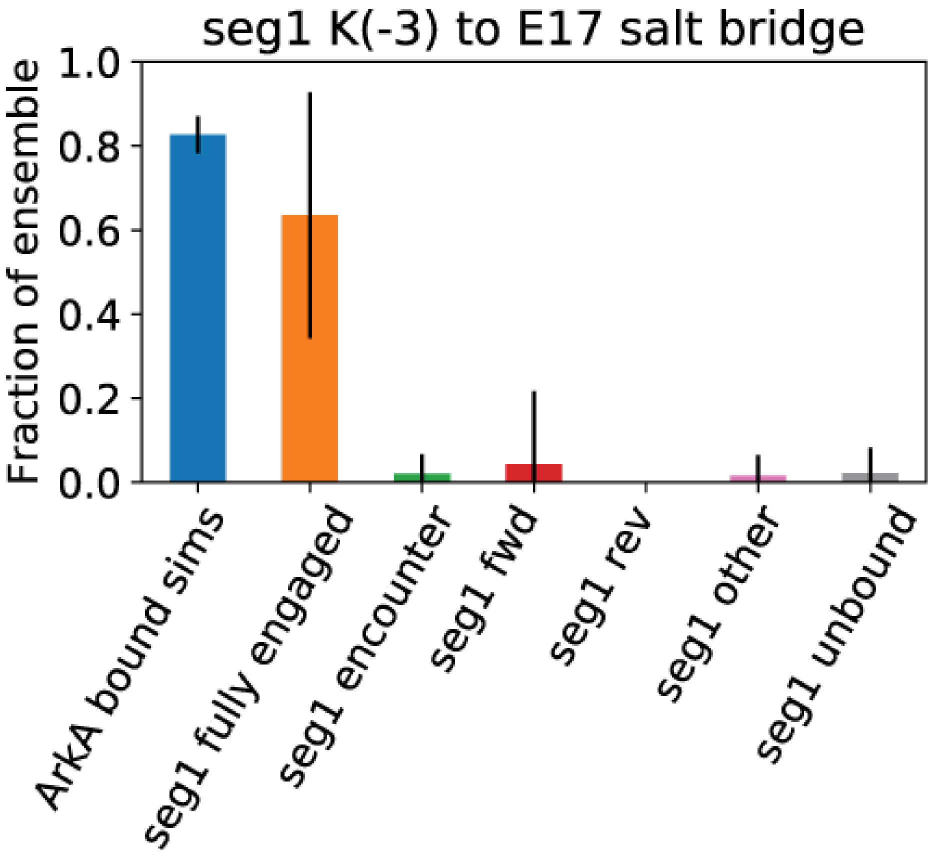


**S19 Fig. Occupancy of the K(-3) to E17 salt bridge for the seg1-AbpSH3 complex in different states.** The first bar on the plot represents the salt bridge occupancy in bound simulations. Error bars represent the standard deviation between independent simulations.

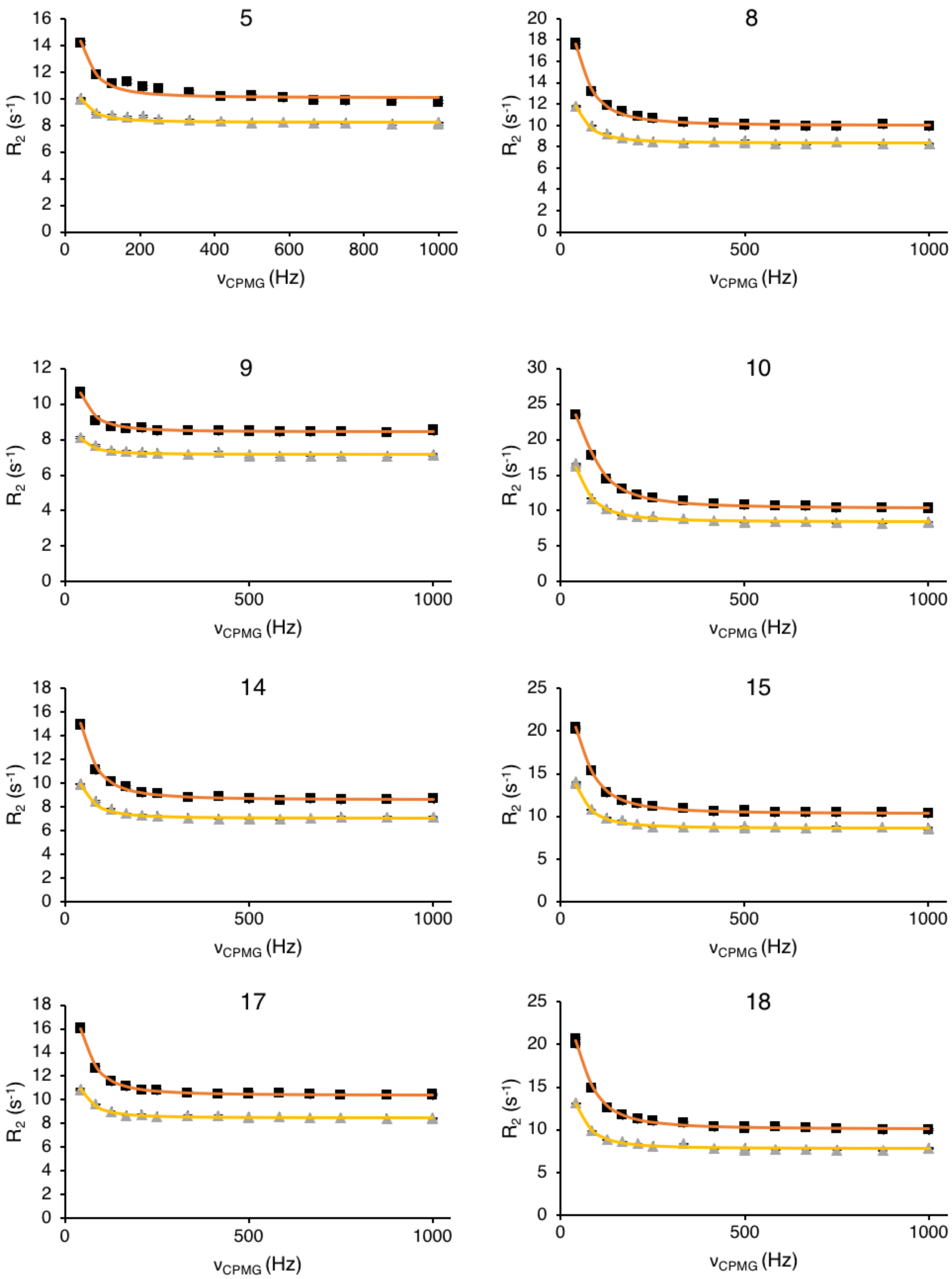
5
8
9
10
14
15
17
18
R2 (s-1)
vCPMG (Hz)
0
200
400
600
800
1000
500

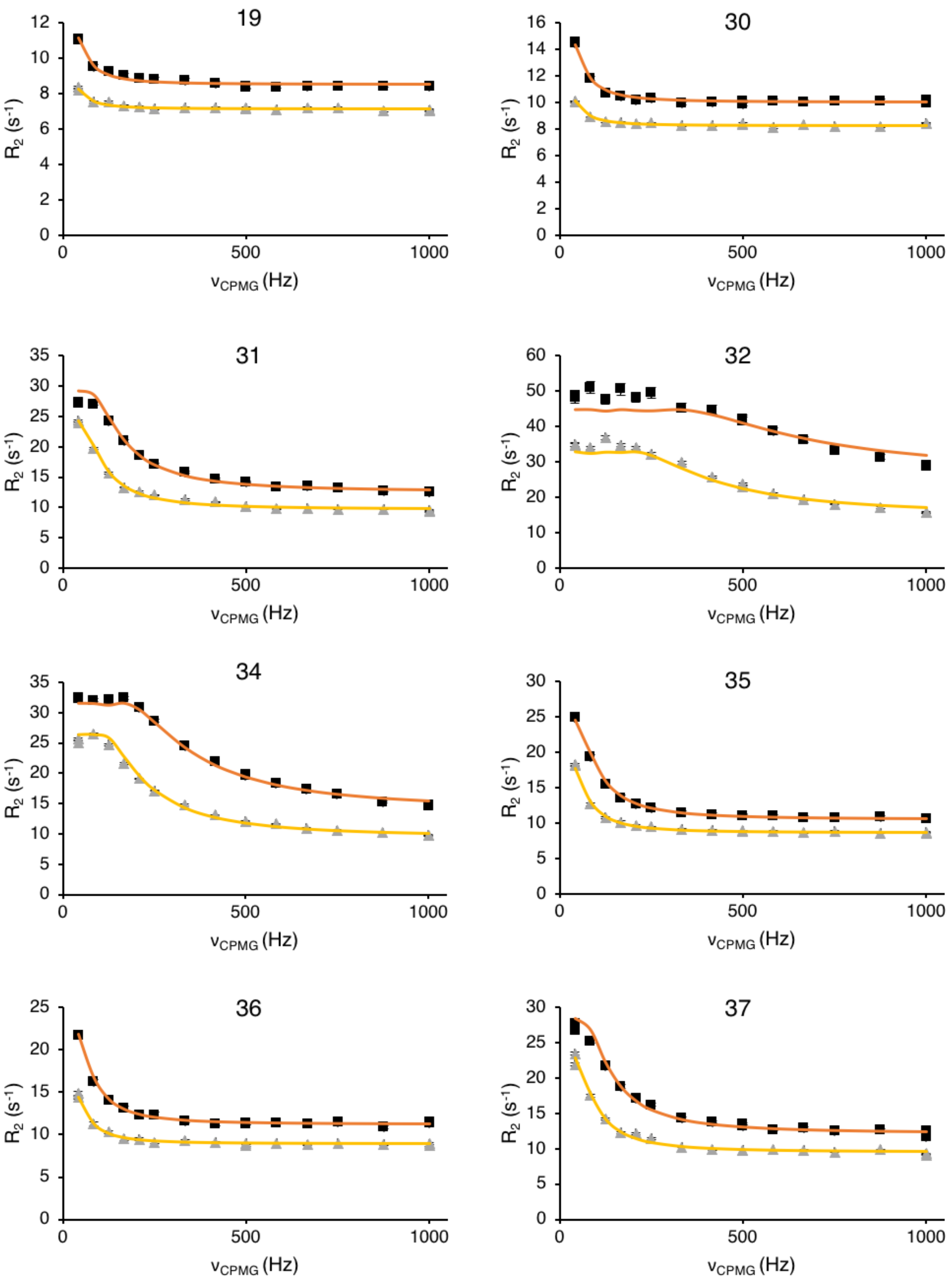
19
30
31
32
34
35
36
37
R2 (s-1)
νCPMG (Hz)

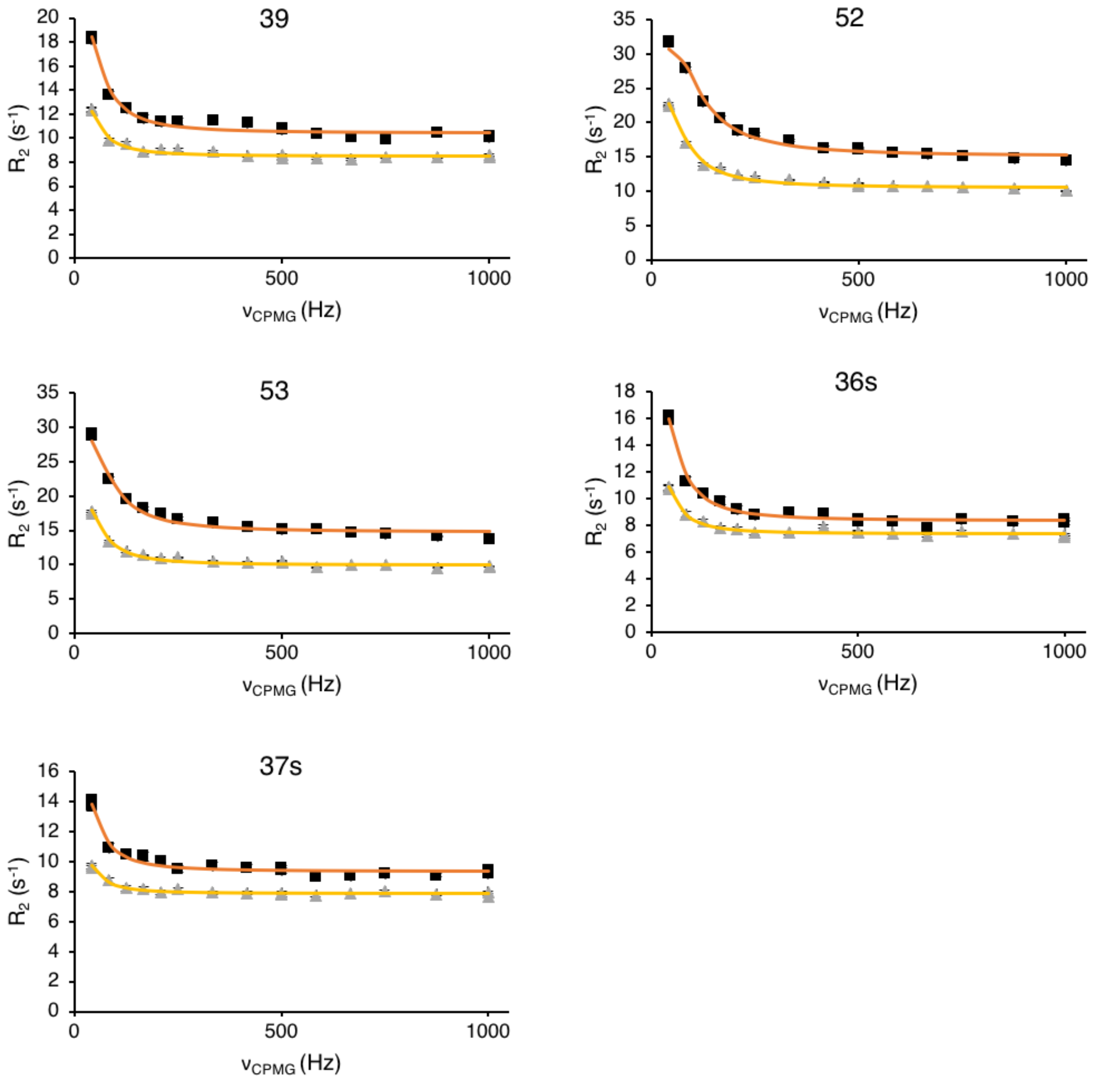


**S20 Fig. NMR $^{15}$N CPMG relaxation dispersion data for the amide signals of 21 AbpSH3 residues at 500 (triangles) and 800 (squares) MHz.** The top of each plot is labeled with the residue number, and 36s and 37s refer to the tryptophan sidechain NH groups.

**S1 Movie. Example simulation of initial ArkA binding to the AbpSH3 domain.**

**S2 Movie. Example simulation of  initial ArkA binding to the AbpSH3 domain.**

**S3 Movie. Simulation of ArkA changing orientation from forward to reverse in the encounter complex.**
