## Supplementary material for "A disordered encounter complex is central to the yeast Abp1p SH3 domain binding pathway": Readme for data

All data is available at https://creativematter.skidmore.edu/faculty_datasets/1

Download and uncompress the files.

The folder system:
There is a folder for the MD running files for each construct which contains: leap, pdb, prmtop, inpcrd, mdin, and scr (batch submission) files 

There is also an analysis folder for each construct, and one folder with all the python scripts used.
